# Oncolytic vaccinia virus expression of a defined peptide-MHCI complex as a precision cancer immunotherapy platform

**DOI:** 10.1101/2024.08.16.608170

**Authors:** S Komant, J Wang, N Favis, C Alex, DH Evans, RS Noyce, TA Baldwin

## Abstract

Oncolytic viruses are immunotherapeutic agents that selectively replicate in and kill tumour cells with the goal of promoting anti-cancer immune responses. Vaccinia virus (VACV) is a strong oncolytic virus candidate as it infects a wide range of cancer cells, is amenable to genetic tailoring and induces potent, long-lasting immunity. Here, we examine the use of genetically engineered oncolytic VACVs (oVACV) to express a variety of MHC-I complexes and the co-stimulatory ligand CD80 to stimulate anti-tumour CD8^+^ T cell responses in two syngeneic cancer models. Tumour antigen specific CD8^+^ T cells were detected in both the tumour and spleen following oVACV treatment, demonstrating the ability to induce tumour targeted T cell responses with viral therapy. oVACV expression of peptide-MHC-I complexes enhanced the efficacy of primary tumour clearance with tumour specific antigen targeting showing the greatest efficacy. Initial tumour clearance following treatment with oVACVs led to variable anti-tumour immune memory depending on the tumour model. Depletion of CD8^+^ T cells prevented therapeutic efficacy of oVACV while combination with αPD-L1 immune checkpoint blockade enhanced tumour clearance. Overall, we demonstrate the ability to generate an oncolytic virus capable of inducing recognition and elimination of tumour cells through VACV-mediated expression of defined peptide-MHC-I complexes.

## Introduction

Cancer immunotherapies rely on harnessing an individual’s immune system to specifically target and eradicate tumours. The ability to induce or augment immunity against cancer cells can be amplified by delivering immune stimulatory molecules systemically or directly to tumours. Oncolytic viruses (OV) are a category of viruses that preferentially infect, replicate within and kill cancer cells. OV-mediated tumour cell killing can also induce anti-tumour immunity. Stimulation of the immune system to provoke anti-tumour immunity classifies these viruses as a form of immunotherapy,^1^ however OVs as single agent virotherapies have struggled to yield the clinical success observed with other forms of immunotherapy.^2–5^ Therefore, developing novel strategies to improve the efficacy of OV therapy and specifically those that generate a potent, long-lasting tumour targeted immune response is needed. Engineering OVs to express defined tumour antigens is a potential method to create such a precision cancer immunotherapy.

The presence of tumour antigens that can provoke an immune response is a central feature of cancer immunotherapy success. Heterogeneity of tumour mutational burdens (TMB) between cancer types limits some tumours in producing neoantigens that can be identified by cytotoxic T lymphocytes (CTL). These genetic and immunological characteristics are used in the categorization of “hot” (immune-inflamed) and “cold” (immune-desolate) tumour microenvironments and correlate with responsiveness to immunotherapy and positive therapeutic outcomes.^6,7^ Immune-inflamed “hot” tumours are characterized by T cell infiltration, elevated TMB and increased PD-L1 expression leading to enhanced response rates to immune check point blockade therapies.^8^ Immunoscore, derived from the density of specific lymphocyte populations in the tumor core and invasive margin, serves as a quantitative prognostic marker for hot tumours and predicting patient outcomes and survival.^9^ High immune scores are strongly correlated with positive prognoses, particularly in colorectal cancers, by quantifying the body’s natural anti-tumor response.^9^ Beyond its predictive value, understanding these immune infiltration patterns is essential for developing and optimizing novel immunotherapies to improve long-term clinical results.^10^ Conversely, tumours with immune-excluded or desolate phenotypes are regarded as “cold” and characterize by confinement of T cells to the tumour periphery or by the complete absence of T cells within the vicinity of the tumour making them less sensitive, or in some cancers, resistant to immunotherapies.^8^ Tumours harbouring major deficiencies in antigen processing and presentation are often found to be immunologically “cold” due to the inability to present cancer neoantigens on their cell surface.^8^ Epigenetic alterations, loss of heterozygosity of human leukocyte antigen (HLA) genes and β2-microglobulin (β2M) loss are common in many cancers and limit the effectiveness of immunotherapies.^11,12^ This phenotype is also observed in murine models of cancer where severe major histocompatibility complex -I (MHC-I) and β2M defects render tumours largely invisible to the anti-tumour effects of T cells. However, even tumours without obvious effects in MHC-I antigen processing and presentation can be classified as “cold” and resistant to immunotherapy. Thus converting “cold” tumours into immunologically “hot” tumours through robust T cell infiltration and recognition of tumour antigens represents a major therapeutic goal.

Vaccinia virus (VACV) is a large, double-stranded DNA virus that has been studied as a promising oncolytic agent.^13^ VACV has numerous endogenous properties that make it a strong cancer therapeutic candidate, including an excellent safety profile due to its use as the agent in the smallpox vaccine campaign,^14^ the capacity to encode large amounts of foreign DNA, easy genetic manipulation, the ability to infect a wide range of cell types, and the ability to induce long lasting and potent cellular immune responses.^15–17^ VACVs have entered clinical studies as a cancer therapeutics^18–20^ with continued research focusing on enhancing the virus’s potential to induce an anti-tumour immune response. Mechanisms explored to improve an anti-tumour immune response include the delivery of immune stimulating molecules like GM-CSF and IL-2.^21,22^ These strategies are tumour antigen agnostic and rely on the immune system “identifying” the correct tumour antigen(s) to target in a sea of self and viral antigens.^23–25^ This antigen agnostic approach is limited by normal self-tolerance mechanisms that may reduce the availability or potency of T cells that can kill tumours. To overcome this limitation, oncolytic viruses have been constructed that express tumour antigens for use as therapeutic cancer vaccines.^26^ A potential barrier in these approaches includes insufficient presentation of all vaccine encoded neoantigens by MHC machinery and pathways as many different tumour types have a reduced capacity to process and present peptide-MHC-I complexes rendering the tumour largely invisible to CD8^+^ T cells.^27–30^ Another recent approach incorporated an oncolytic VACV (oVACV) that expressed a target for CAR-T cells with infusion of appropriate CAR-T cells to focus the immune response on the tumour.^31^ Flagging the tumour cells for recognition by CAR-T cells by using oVACV to express a CAR ligand overcomes the limitation of pMHC-I down-regulation but introduces the need to create CAR-T cells, requires high levels of tumour infection and reduces the potential to generate anti-tumour T cell memory responses. Engineering an oncolytic VACV that expresses a pre-loaded pMHC-I complex presenting a defined tumour peptide may create an optimum therapeutic cancer vaccine, as it bypasses the need to have tumour antigen processed and presented, leading to responses against the specific tumour antigens being introduced.

Here, we created and tested an engineered oVACV encoding an exogenous peptide-MHC-I (pMHC-I) gene expressed as a single chain trimer (SCT) complex with β2-microglobulin (β2M)^32–34^ in two separate cancer models. Specifically, we determined that expression of the pMHC-I complex by oVACV induced an antigen-specific T cell response and enhanced anti-tumour immunity resulting in tumour clearance in some cases. The engineered oVACV required CD8^+^ T cells for efficacy and showed additive effects with αPD-L1 checkpoint blockade. Strong immune memory responses were observed when animals were treated with the oVAC expressing relevant pMHC-I. Collectively, this research supports the ability to bolster an anti-tumour specific immune response using modified VACV as both an oncolytic agent and a vector for immunotherapeutic antigen delivery and confirms the validity of VACV as a precision cancer immunotherapy.

## Results

### Generation of engineered oVACV variants

We previously examined the impact of deleting specific VACV genes (alone or in combination) in the context of cancer virotherapy. A combination of deletions that improved responses to two different breast cancer cell lines were identified.^35^ This VACV strain, VACVΔ3X, contains three gene deletions, *J2R*, *B8R* and *B18R*. *J2R* encodes a viral thymidine kinase analog that when removed renders the virus tumour selective.^36,37^ *B8R* and *B18R* regulate the host type I and II interferon (IFN) pathways respectively.^38,39^ VACVΔ4x was generated from the VACVΔ3X backbone by deleting the *M2L* locus; a viral protein shown to bind the CD80 costimulatory molecule and interfere with T cell activation.^40^ CD80 was introduced into the VACVΔ4x *B8R* locus to enhance co-stimulation by virally infected cells (Figure 1A). Finally, to drive an antigen specific anti-tumour response we genetically engineered VACVΔ4x-CD80 to express a covalently linked single chain trimer (SCT)^33,34^ or single chain dimer (SCD) composed of the major histocompatibility complex-I heavy chain, β2-microglobulin, with or without a defined peptide, respectively, in the *M2L* locus (Figure 1A). For proof-of-concept experiments, one iteration of the SCT encoded the SIINFEKL peptide from chicken ovalbumin (OVA_(257-264)_). The OVA_(257-264)_ peptide (OVAp) was selected as a well-studied foreign antigen used frequently in murine studies.^41,42^ Additionally, the tumour associated peptide Trp2_(180-188)_, was selected for use with the B16F10 melanoma model as a tumour associated antigen, as it is from an endogenously expressed melanin protein and has been previously used as a therapeutic cancer vaccine target.^43,44^ The SCT designed for the EMT6 murine breast cancer model encodes a previously identified mutated self-peptide that can induce a CD8^+^ T cell response and not cross react with the wildtype protein to represent a tumour specific antigen.^45^ These mutations and immune modulatory insertions allow for modified oVACV tumour selectivity and potential to stimulate an anti-tumour immune response.

**Figure 1.**
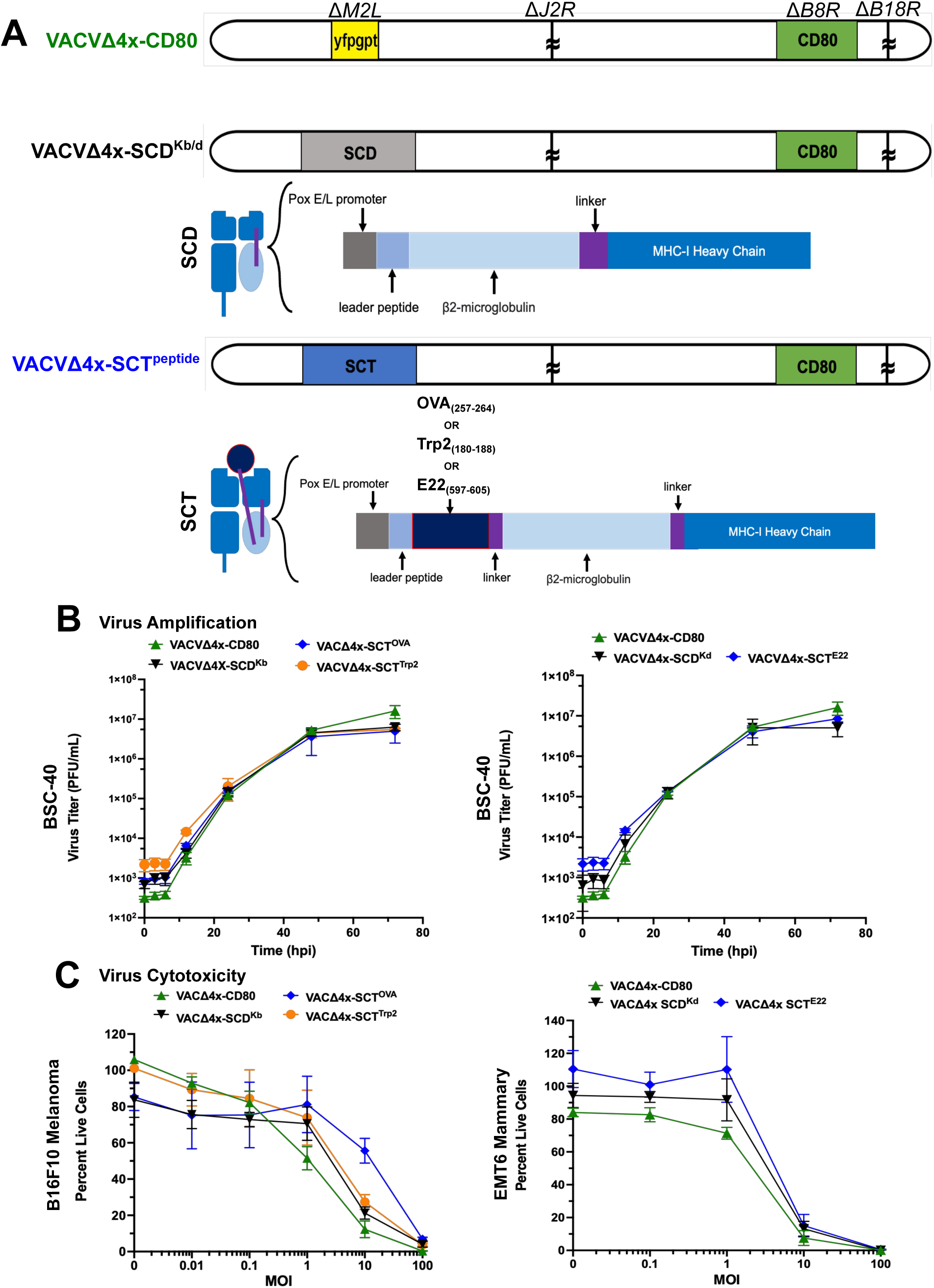
Oncolytic VACVs and in vitro growth properties. A, Genome maps of oVACV constructs rendered attenuated through four deletions in the VACV Western Reserve backbone at the M2L, J2R, B8R and B18R loci. At each deletion site there is an insertion of either yellow fluorescent protein-guanine phosphoribosyl transferase (yfp-gpt), CD80, single chain dimer (SCD) or single chain trimer (SCT), where the SCT is expressing one of three peptides, OVA_(257-264)_, Trp2_(180-188)_, E22_(597-506)_. B, Viral growth kinetics of oVACVs on BSC-40, immortalized African green monkey kidney cells. C, Almar blue cytotoxicity assay 72 hours post infection in B16F10 and EMT6 tumour cells. Error bars denote standard deviation (SD) from 3 independent experiments.

### Growth and cytotoxicity properties of engineered oVACV *in vitro*

Viral growth rates and cytolytic capacity are important characteristics of viral fitness and oncolytic potential. Following genetic engineering of the various recombinant oVACVs, we first evaluated their growth properties. BSC-40 cells were infected at a MOI of 0.03 and viral titres were measured at various time points over 72 hours (Figure 1B). All virus variants exhibited similar growth patterns over time when plated at equal MOIs, indicating that the genetic modifications did not impact viral fitness.

The oVACV variants were also evaluated for tumour cell killing capacity *in vitro* using a cytotoxicity assay where tumour cells were infected over 72 hours and metabolism was assessed as a marker of cell viability after infection. All virus variants were similarly cytotoxic in both B16F10 and EMT6 cell lines (Figure 1C) except for the VACVΔ4x-SCT^OVA^ virus at a MOI of 10 in B16F10 cells, however this was not significant (Figure 1C). Together these data show that deleting the described immunomodulatory genes and inserting immunostimulatory protein producing genes had minimal effect on oVACV growth or cytotoxicity *in vitro*.

### Expression of SCT and SCD activates CD8^+^ T cells *ex vivo*

We next evaluated the cell surface expression of virally encoded CD80, SCT or SCD proteins individually or in combination on B16F10 cells following infection. B16F10 melanoma cells have well-described impairments in the expression of various components of the MHC-I antigen processing and presentation pathway resulting in severely limited pMHC-I cell surface expression.^46,47^ B16F10 tumour cells were infected with the VACVΔ4x-SCT^OVA^ virus *in vitro* and 24h post-infection, we stained the cells with anti-CD80 and the 25D1.16 antibody that specifically recognizes K^b^-OVAp complexes. Expression of both CD80 and K^b^-OVAp indicated that expression of the pre-formed SCT was not impacted by intrinsic impairments in MHC-I antigen processing and presentation (Figure 2A). To determine if the virally expressed SCT^OVA^ complex induced CD8^+^ T cell activation, oVACV infected B16F10 tumour cells and whole CFSE labelled splenocytes from OVA_(257-64)_ specific OT-I TCR transgenic mice were co-cultured for 72 hours. T cell surface expression of CD25 and CD69 were assessed as indicators of T cell activation.

**Figure 2.**
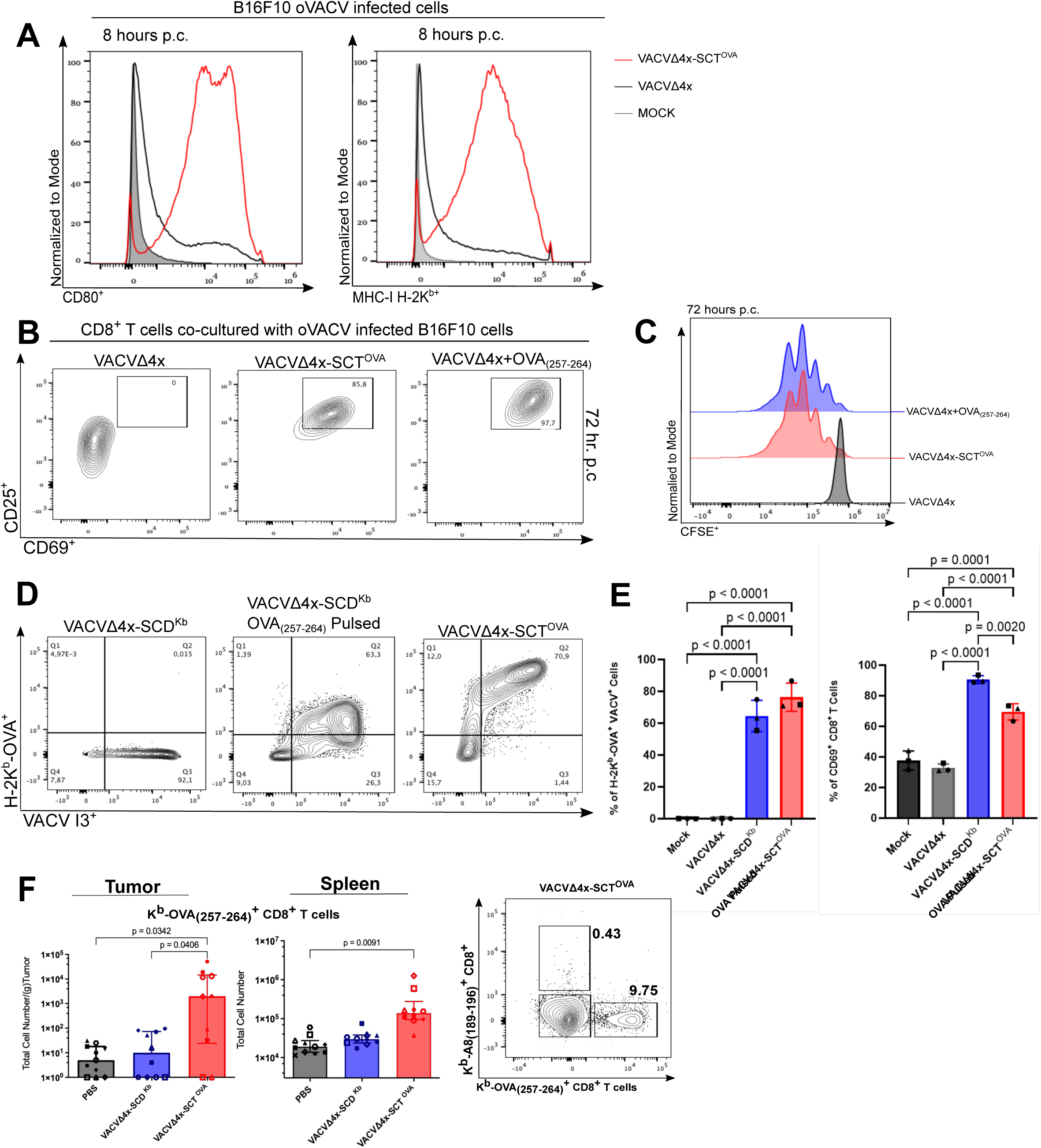
Tumour cell surface expression of the MHC-I SCT complex and CD80 activates antigen specific T cells *in vitro* and expands OVA_(257-264)_ specific T cells *in vivo*. A, VACVΔ4x-SCT^OVA^ infected tumour cells were stained with anti-CD80 and anti-H-2K^b^-OVA antibodies. Representative flow cytometry plots from three independent experiments are shown. B, Flow cytometry plots of OT-I T cells cultured with VACVΔ4x-SCT^OVA^ infected B16F10 tumour cells stained with anti-CD69 and anti-CD25 antibodies. Representative flow cytometry plots from three independent experiments are shown. C, Flow cytometry plots of CFSE labelled OT-I cells following 72 hour co-culture with VACVΔ4x-SCT^OVA^ infected B16F10 cells. A representative CFSE plot shown from three independent experiments. D, Cell surface expression of H-2K^b^-OVA on OVA_(257-264)_ pulsed B16F10 tumour cells infected with oVACV variants. Representative flow cytometry plots from three independent experiments are shown. E, H-2K^b^-OVA complex expression on oVACV infected cells, and CD69 surface expression on OT-I T cells co-cultured with OVA_(257-264)_ pulsed and oVACV infected B16F10 cells overnight from three independent experiments. F, OVA_(257-264)_ specific CD8^+^ T cells from B16F10 tumour and spleen were identified with OVA-K^b^ tetramers following VACVΔ4x-SCT^OVA^ treatment mice 14 days after the first virus treatment. A representative flow cytometry plot from two independent experiments is shown (n=10). Statistical analysis was preformed using one-way ANOVA from up to three independent trials, where p<0.05 is significant and error bars denote mean (SD) or medians and inter-percentile range for data represented on a logarithmic scale.

VACVΔ4x-SCT^OVA^ infection of B16F10 cells led to activation marker expression on most OT-I T cells at 72 hours post co-culture *ex vivo* (Figure 2B). This activation closely matched the activation of OT-I splenocytes stimulated with soluble OVA_(257-64)_ peptide in the presence of VACVΔ4x infected B16F10 cells (Figure 2B). Further, these activated CD8^+^ T cells underwent numerous rounds of cell division, with VACVΔ4x-SCT^OVA^ infected cells inducing proliferation similar to that of the peptide stimulated OT-I T cells (Figure 2C).

We additionally evaluated the ability of the virally expressed SCD, lacking a covalently linked peptide, to be loaded with exogenous peptide. B16F10 tumour cells were infected with VACVΔ4x-SCD^Kb^ overnight followed by pulsing with OVA_(257-64)_ peptide for 1 hour. After infection with VACVΔ4x-SCD^Kb^ and peptide pulsing, approximately 60% of tumour cells expressed H-2K^b^-OVAp on the cell surface (Figure 2D and E), and the co-culture of peptide pulsed VACVΔ4x-SCD^Kb^ infected B16F10 cells with OT-I splenocytes induced CD69^+^ expression on over 80% of CD8^+^ T cells (Figure 2E). We then evaluated the ability to generate an OVA_(257-264)_ specific CD8^+^ T cell response in tumour bearing animals following intratumoural oVACV treatment. B16F10 tumours were implanted orthotopically in the flanks of male C57BL/6 mice. Palpable tumours were treated with 3 injections of oVACV at 1.0×10^7^ PFU 48 hours apart. On day 14 post-treatment, tumours were harvested for immune analysis. A 2-log increase in the number of OVA_(257-264)_ specific CD8^+^ T cells was found in the tumours of mice treated with VACVΔ4x-SCT^OVA^ compared to all other treatment groups (Figure 2F). OVA_(257-264)_ specific CD8^+^ T cells were also detected in the spleens of mice treated with VACVΔ4x-SCT^OVA^ where the number of OVA_(257-264)_ specific CD8^+^ T cells was significantly increased compared to PBS (Figure 2F). Collectively, these data demonstrated expression of the SCT complex and CD80 on B16F10 tumour cells following infection with VACVΔ4x-SCT^OVA^ resulted in activation and proliferation of antigen specific CD8^+^ T cells *in vitro* and *in vivo*. Additionally, infection with VACVΔ4x-SCD^Kb^ and peptide pulsing can lead to antigen loading of the empty MHC-I complex and induce CD8^+^ T cell activation.

### Administration of engineered oVACV to B16F10 melanomas induces changes to the intratumoral immune landscape

The ability to induce T cell activation and proliferation *in vitro* and *in vivo* through cell surface expression of the SCT^OVA^ complex and CD80 led us to investigate if the treatment of tumours *in vivo* using an SCT expressing a relevant tumour associated antigen (TAA) could result in the induction of a tumour specific immune response. Here, we again used the B16F10 murine melanoma model and a VACVΔ4x expressing no exogenous pMHC-I, an empty H-2K^b^ SCD or a SCT encoding H-2K^b^ and the TAA Trp2_(180-188)_. The impaired endogenous pMHC-I expression within the B16F10 model provides an opportunity to specifically investigate the impact of virally expressed SCT or SCD MHC-I complexes on immune responses. We monitored immune activation, tumour regression and animal survival after virus treatment with the engineered oVACV variants.

Tumours were treated as described in Figure 3A, where a cohort of animals was used for immune cell analysis and a second cohort of animals was used to monitor tumour volume over time. Oncolytic virus treatment altered the proportion and number of immune cells recovered from the tumour (Figure 3B and C). Generally, there was an overall trending increase in the proportion of infiltrating CD8^+^ T cells and a trending decrease in the proportion of B cells in the virus treated groups (Figure 3B). An increase in the total number of tumour infiltrating CD4^+^ and CD8^+^ T cells was observed with VACVΔ4x-SCT^Trp2^ treatment compared to PBS and VACVΔ4x-CD80 treated groups (Figure 3D and E). When analyzing suppressive T cells, known as regulatory T cells (Tregs) categorized as being CD4^+^, CD25^+^, and FOXP3^+^, there was a significant increase in the total number of Tregs infiltrating the tumour following VACVΔ4x-SCT^Trp2^ treatment (Figure 3E). However, when the frequency of regulatory T cells of total CD4^+^ T cells was evaluated, there was a significant decrease in the frequency of regulatory T cells in all oVACV treated groups compared to PBS (Figure 3E). In the non-T cell compartment, a trend toward increasing proportions of NK1.1^+^ (CD4^−^ CD8^−^ CD19^−^) natural killer-like cells was observed following VACVΔ4x-SCT^Trp2^ treatment (Figure 3B), though this did not lead to a significant difference in the total number of NK1.1^+^ NK-like cells between groups (Supplemental Figure 2A). NK1.1^+^ CD8^+^ T cells have been identified as potent mediators of the anti-tumour response,^48,49^ and we observed a trending increase in the number of NK1.1^+^ CD8^+^ T cells when the tumour was oVACV treated, although they were observed at a lower frequency of total CD8^+^ T cells following VACVΔ4x-SCT^Trp2^ treatment (Figure 3F). Expression of PD-1 on the T cell surface is an early activation marker, and with prolonged expression may also indicate a functional transition toward T cell exhaustion.^50,51^ We found an increase in the number and frequency of programmed cell death protein 1 (PD-1) expressing CD8^+^ T cells in the VACVΔ4x-SCT^Trp2^ treated group compared to PBS and VACVΔ4x-CD80 (Figure 3F).

**Figure 3.**
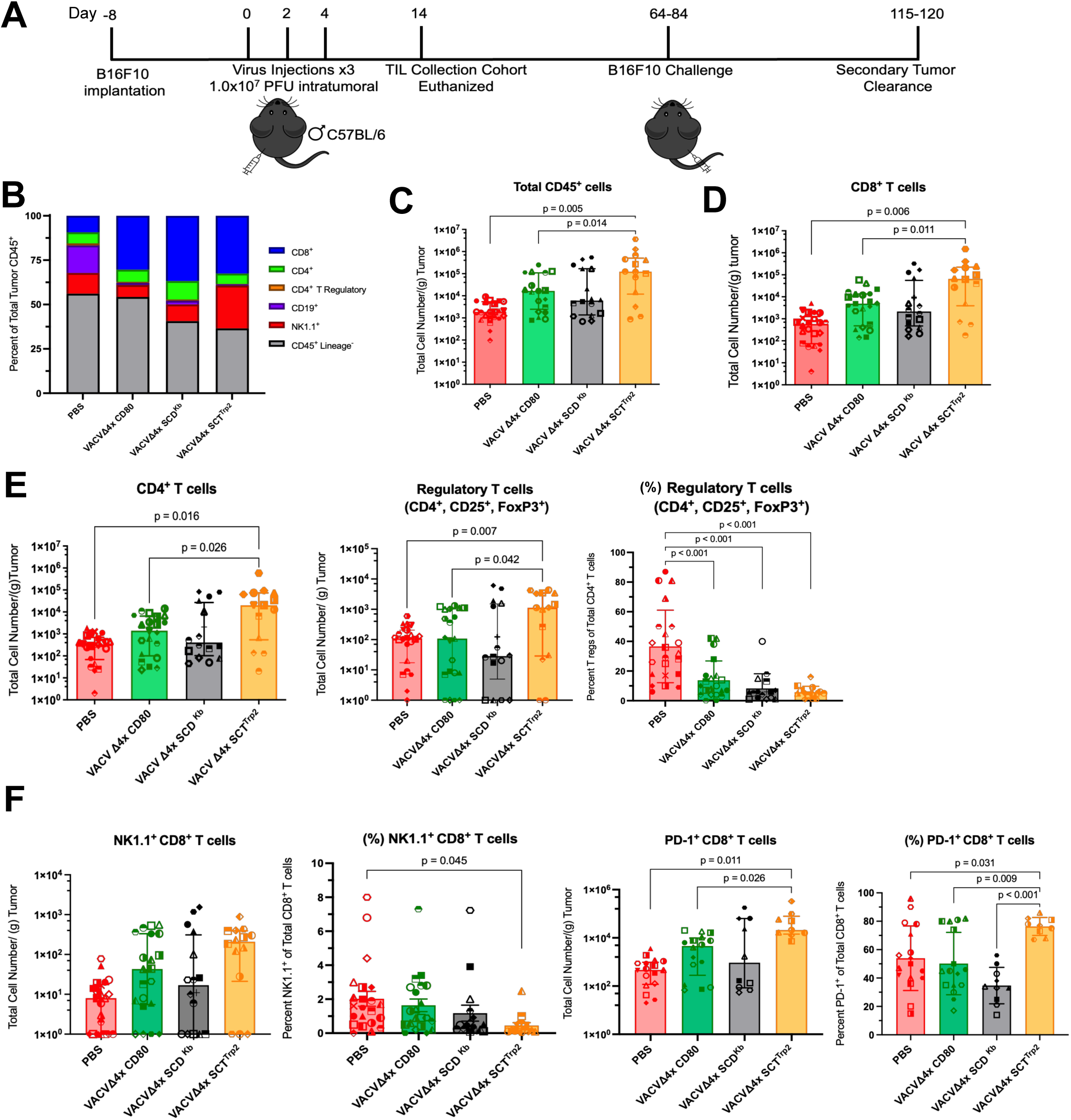
Oncolytic VACV induces altered immune infiltration and cell surface marker expression in B16F10 tumours. A, Experimental design to assess survival, tumour growth and tumour infiltrating lymphocyte population changes after treatment with oVACVs. Six to eight-week-old male C57B/6 mice were injected with B16F10 tumour cells subcutaneously in the flank. Once palpable, three intratumoural 1.0×10^7^ PFU doses of oVACVs were administered. A cohort of mice were euthanized 14 days after the first virus dose, where tumour and spleen were collected for immune analysis. B-C, Total lymphocytes (CD45^+^) and the immune populations infiltrating the tumour in PBS and oVACV treated mice. D, Total CD8^+^ T cells infiltrating the tumour in PBS and oVACV treated mice. E, Total CD4^+^ T cells and regulatory T cells infiltrating the tumour, and the frequency of regulatory T cells of total CD4^+^ T cells in PBS and oVACV treated mice. F, Total CD8^+^ T cells expressing NK1.1 or PD-1 and the frequency of NK1.1^+^ or PD-1^+^ CD8^+^ T cells of total CD8^+^ T cells in PBS and oVACV treated mice. Statistical analyses were performed using one-way ANOVA from three independent trials and error bars denote mean (SD) or medians and inter-percentile range for data represented on a logarithmic scale.

### Administration of engineered oVACV expressing a TAA induced tumour antigen specific T cell responses

In addition to examining bulk CD8^+^ T cell populations, we used H-2K^b^ tetramers loaded with Trp2_(180-188)_ to measure the Trp2_(180-188)_ specific CD8^+^ T cell response in treated tumours (Figure 4A). We observed a significant increase in the number of Trp2_(180-188)_ specific CD8^+^ T cells infiltrating tumours following treatment with VACVΔ4x-SCT^Trp2^ compared to all other groups (Figure 4B), demonstrating the ability of oVACV treatment to expand the Trp2_(180-188)_ specific CD8^+^ T cell population infiltrating the tumour. Similarly, spleens of treated mice contained increased numbers of Trp2_(180-188)_ specific CD8^+^ T cells in mice treated with VACVΔ4x-SCT^Trp2^ compared to all other treatments (Figure 4C and D). Further, we evaluated the expansion of the VACV specific CD8^+^ T cells in both the tumour and spleen. Here we used H-2K^b^ tetramers loaded with VACV A8_(189-196)_, the immunodominant H-2K^b^ peptide of our modified oVACV, to detect virus specific CD8^+^ T cells (Figure 4B and D). oVACV treatment led to an increase in the total number of VACV A8_(189-196)_ specific CD8^+^ T cells both in the tumour and spleen compared to PBS, though this was not statistically significant (Figure 4B and D).

**Figure 4.**
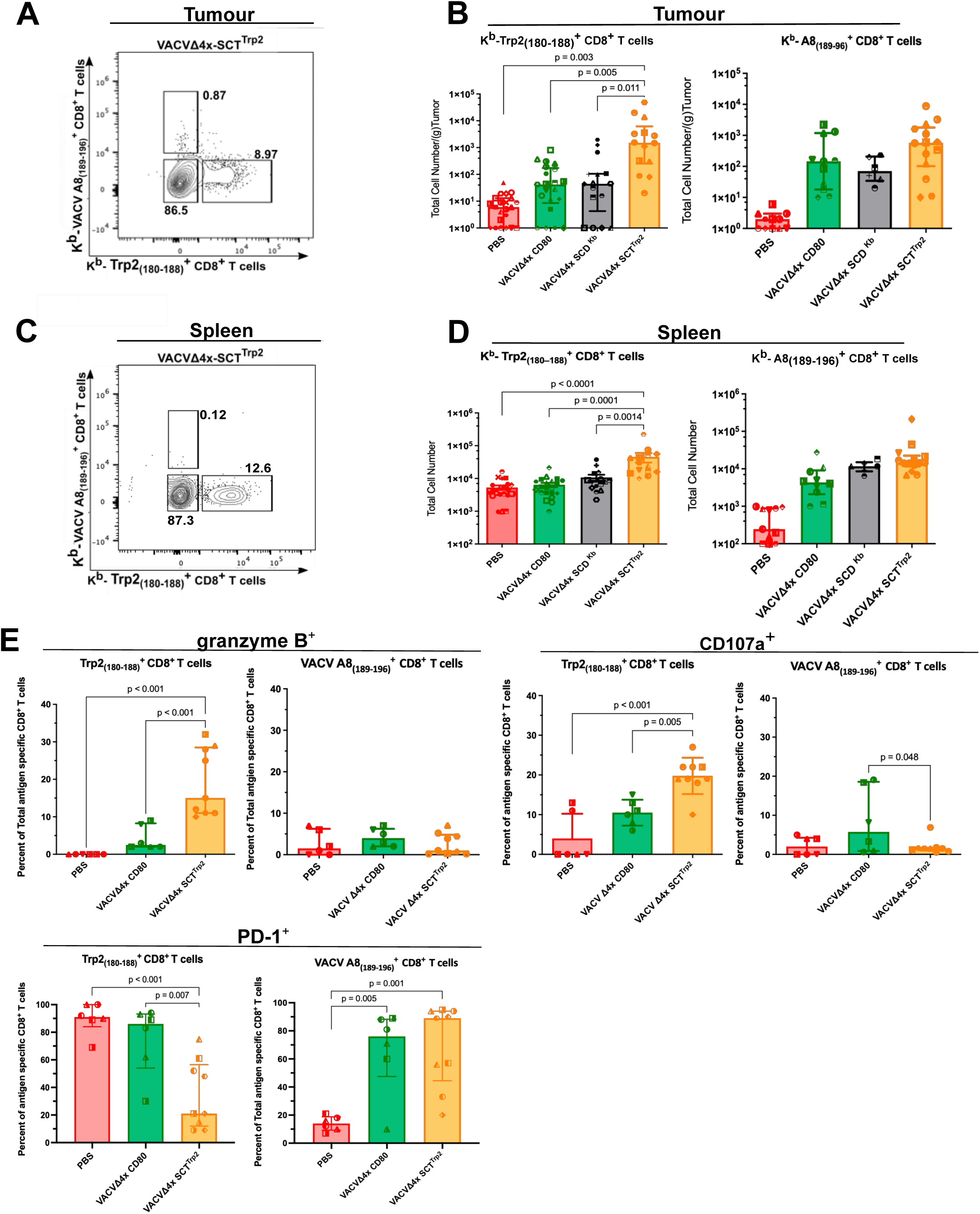
Antigen specific CD8^+^ T cells infiltrate B16F10 tumours after oncolytic VACV treatment and differentially express markers of T cell effector potential. A, Trp2_(180-188)_ and VACV A8_(189-196)_ antigen specific CD8^+^ T cell numbers infiltrating tumours from PBS and oVACV treated mice. B, A representative flow cytometry plot for data in A is shown. C, Trp2_(180-188)_ and VACV A8_(189-196)_ antigen specific CD8^+^ T cells populations in the spleen from PBS and oVACV treated mice. D, A representative flow cytometry plot for data in C is shown. E, Frequency of tumour infiltrating antigen specific CD8^+^ T cells (Trp2_(180–188)_^+^ VACV A8_(189-196)_^+^) expressing intracellular granzyme B, cell surface CD107a or PD-1 in PBS and oVACV treated mice. Statistical analyses were performed using one-way ANOVA from three independent trials (n>8 mice) where p<0.05 is significant and error bars denote mean (SD) or medians and inter-percentile range for data represented on a logarithmic scale.

To mediate tumour regression these antigen specific CD8^+^ T cells need to be functional effectors. One mechanism of CD8^+^ T cell effector function is the production of cytotoxic molecules and subsequent degranulation. Intracellular granzyme B, an apoptosis inducing protease, and cell surface CD107a, a lysosomal protein that becomes exposed on the cell surface after CD8^+^ T cell degranulation, were examined in Trp2_(180-188)_^+^ CD8^+^ T cells to determine if these tumour antigen specific cells may also have effector potential. An increase in the frequency of Trp2_(180-188)_ specific CD8^+^ T cells that contained intracellular granzyme B or expressed cell surface CD107a was observed in VACVΔ4x-SCT^Trp2^ treated mice compared to PBS and a non-MHC-I expressing virus control (Figure 4E). However, these changes were not observed with CD8^+^ T cells specific for the VACV antigen A8_(189-196)_ (Figure 4E). In addition, PD-1 expression was further examined on antigen specific CD8^+^ T cells. In VACVΔ4x-SCT^Trp2^ treated mice, there was a significant decrease in the frequency of Trp2_(180-188)_ specific CD8^+^ T cells expressing PD-1 compared to control groups (Figure 4E). In contrast, VACV A8_(189-196)_ specific CD8^+^ T cells infiltrating the tumours of VACVΔ4x-SCT^Trp2^ treated mice, were significantly increase in PD-1 expression compared to controls (Figure 4E). Together these data indicate the ability of oVACV expressed SCT to expand the number of tumour antigen specific CD8^+^ T cells in the tumour and spleen of treated mice and generate tumour antigen specific effector T cells.

### Administration of engineered oVACV improves B16F10 melanoma tumour clearance in a CD8 T cell dependent manner and is enhanced by anti-PD-L1 checkpoint blockade

We also analyzed the ability of oVACV treatment to impact tumour growth and enhance animal survival. All mice treated with PBS reached tumour burden endpoint by day 20 after the first virus treatment, while many mice treated with the oVACV variants reached tumour burden endpoint after day 20 or in some cases cleared the tumour (Figure 5A). While little difference in the ability to induce tumour clearance was observed between the VACVΔ4x-SCT^Trp2^ and VACVΔ4x-SCD^Kb^ groups, mice treated with VACVΔ4x-CD80 showed a higher survival rate and greater tumour clearance (Figure 5A and Supplemental Figure 3A) although this difference was not statistically significant. In mice where tumours regressed and were no longer palpable, tumour clearance was verified by following animals for at least 30 days after clearance was first determined. VACVΔ4x-CD80 treatment yielded complete responses (CR) in 16 out of 25 mice (64%) (Figure 5A), while treatment with VACVΔ4x-SCD^Kb^ and VACVΔ4x-SCT^Trp2^ produced 9 out of 24 (37.5%) and 17 out of 37 (45.9%) CRs, respectively (Figure 5A). Overall, treatment with any of the recombinant oVACVs was superior to PBS.

**Figure 5.**
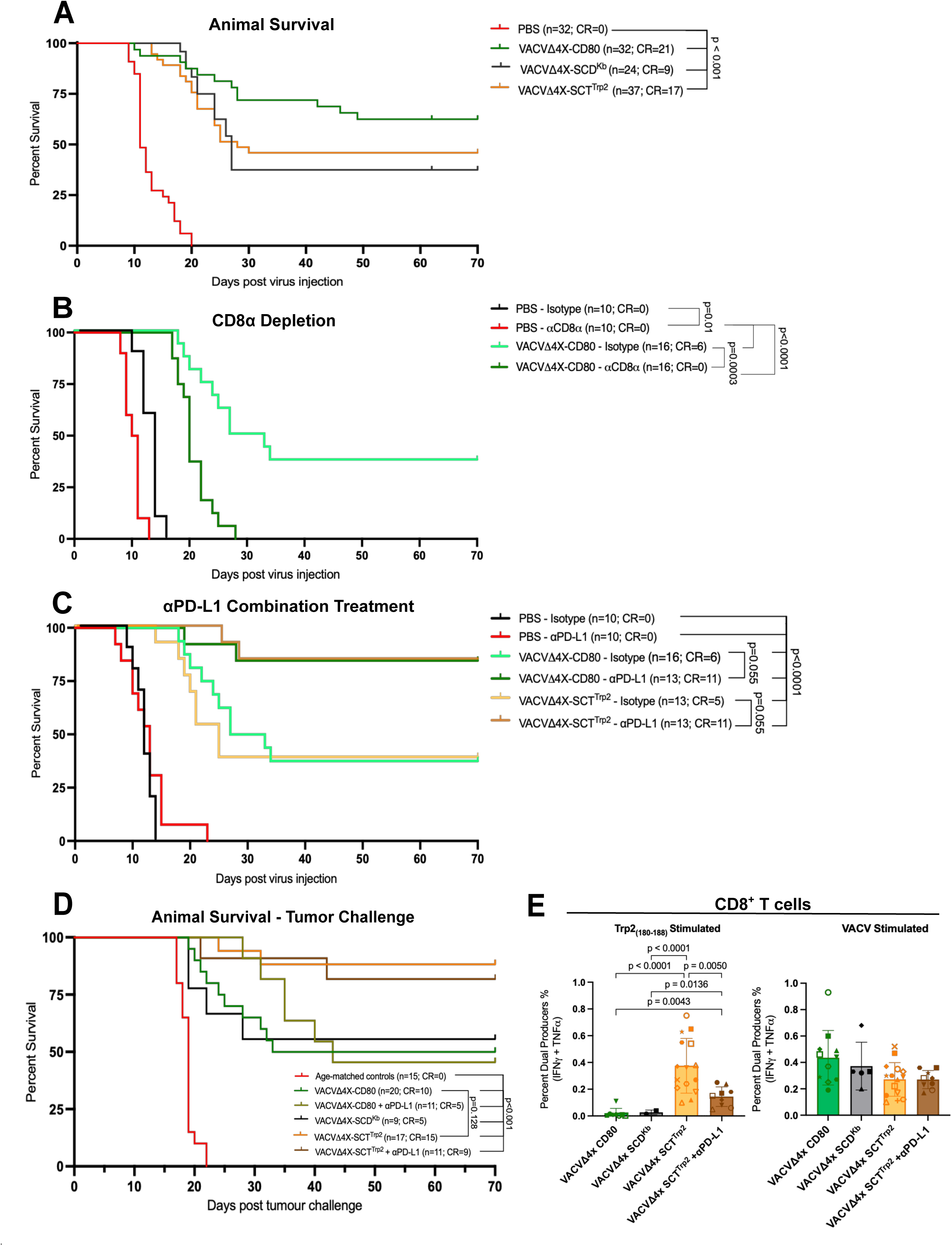
Oncolytic VACVs altered animal survival and potential therapeutic outcomes in B16F10 tumour bearing mice. A, Kaplan-Meier plot of animal survival after intratumoral treatment with PBS or oVACVs. B, Kaplan-Meier plot of animal survival after intratumoral treatment with PBS or oVACVs and CD8^+^ T cell depletion. C, Kaplan-Meier plot of animal survival after intratumoral PBS or oVACVs and αPD-L1 combination therapy. D, Kaplan-Meier plot of animal survival after secondary tumour implantation in the opposite flank of primary tumour cleared mice. E, CD8^+^ T cells from mice that cleared primary and secondary tumour produce both IFN*γ* and TNF*⍺* following *ex vivo* peptide stimulation in the presence of Brefeldin A for 4 hours. Statistical analyses were performed using one way ANOVA to compare between groups at day 70 from up to six independent experiments, where p<0.05 is significant and error bars denote mean (SD). The represented isotype group data in B-C are the same.

To determine the extent to which CD8^+^ T cells play a role in tumour clearance in the B16F10 model treated with VACVΔ4x-CD80, we depleted CD8^+^ T cells prior to oVACV administration. Following CD8^+^ T cell depletion, no mice treated with VACVΔ4x-CD80 cleared their tumour (Figure 5B) although treatment with oVACV and CD8^+^ T cell depletion extended survival over animals treated with PBS and CD8^+^ T cell depletion. In addition to the importance of CD8^+^ T cells, several studies have shown benefits of treating tumours with OVs and immune checkpoint blockade therapy (ICBT) in combination.^52,53^ This combination therapy has also been evaluated as a treatment strategy for advanced melanomas in a phase III trial.^54^ Similarly, we observed an increase in animal survival with combination VACVΔ4x-SCT^Trp2^ and αPD-L1 ICBT (11 out of 13 CRs) which was identical to VACVΔ4x-CD80 and αPD-L1 ICBT (11 out of 13 CRs) and was greater than either virus alone (Figure 5C). As previously reported, αPD-L1 ICBT alone is largely ineffective in treating B16F10 melanoma.^55,56^

Finally, as treatment with oVACV variants led to the clearance of tumour in some mice, we investigated whether long lasting anti-tumour immunity was established in the cured mice. Mice that cleared their primary tumour were reimplanted orthotopically with 1.0×10^5^ B16F10 tumour cells in the opposite and paralleled flank 30 days after the last mouse cleared their primary tumour (Figure 3A). Age-matched, naïve control mice were similarly implanted with tumour cells. All aged-matched control mice reached tumour burden endpoint just after day 20 as seen previously (Figure 5D). While VACVΔ4x-CD80 treated mice resulted in the largest frequency of complete responses following primary tumour treatment, only 10 out of 20 (50%) mice cleared the secondary tumour challenge (Figure 5D). Similarly, although αPD-L1 ICBT enhanced primary tumour clearance when combined with VACVΔ4x-CD80 treatment, the clearance of secondary tumours was limited to 5 out of 11 (45%) of mice (Figure 5D). The ability to reject secondary implanted cells was significantly increased in mice originally treated with VACVΔ4x-SCT^Trp2^ alone or in combination with αPD-L1 ICBT where 15 out of 17 (88%) and 9 out of 11 (82%) mice exhibited complete responses, respectively (Figure 5D). This indicates the potential to generate robust immunological memory responses when an oVACV expressing a pMHC-I complex presenting a TAA is used as a primary therapy. Since we observed an increase in granzyme and CD107a following primary rumour treatment we wanted to examine the functionality of CD8^+^ T cells post 2° tumour rejection. Splenocytes from all mice that rejected the 2° tumour were stimulated with either Trp2_(180-188)_ or a pool of four VACV peptides *ex vivo* and the production of IFNγ and TNFα in CD8^+^ T cells was measured. Trp2_(180-188)_ peptide stimulation resulted in a significantly greater frequency of CD8^+^ T cells that produced both cytokines in mice initially treated with VACVΔ4x-SCT^Trp2^ with or without αPD-L1 ICBT (Figure 5E). In contrast, animals initially treated with VACVΔ4x-CD80 or VACVΔ4x-SCD^Kd^ that then cleared 2° tumour did not show a robust Trp2_(180-188)_ specific response (Figure 5E). Splenic CD8^+^ T cells from all tumour immune mice also produced IFNγ and TNFα in response to the pool of VACV peptides regardless of the oVACV variant used for treatment. Overall, these data validate the ability to stimulate tumour antigen specific CD8^+^ T cell responses that are not only capable of inducing tumour cell regression *in vivo* but further results in the generation of robust immunological memory responses that leads to tumour protection upon challenge.

### Engineered oVACVs reshape the tumour immune landscape and induce tumour specific CD8^+^ T cells in EMT6 mammary tumours

Having established the ability of VACV-expressed pMHC-I to induce antigen-specific CD8^+^ T cell responses to foreign and tumour associated antigens when tumours were impaired in endogenous MHC-I presentation, we transitioned to investigate the immunostimulatory oVACV variants in a model with intact antigen processing and presentation machinery. We used the EMT6 model, a mammary carcinoma cell line derived from a spontaneous hyperplastic alveolar nodule of a BALB/c mouse and is considered representative of the triple negative breast cancer model.^57^ EMT6 cells are slower growing, weakly immunogenic and can spontaneously metastasize to distant organs.^58^ Furthermore, we selected a recently identified tumour specific antigen (TSA), E22_(597-605)_ peptide to be presented in the SCT complex.^45^ EMT6 cells have a single amino acid substitution in the native protein which created an immunogenic peptide that can be recognized by CD8^+^ T cells in BALB/c mice.

Orthotopic mammary tumours were treated as described in Figure 6A. For the EMT6 model, tumour establishment occurred over 10 days, and examination of the immune composition and response in the tumour and spleen was examined 14 days after the first virus injection. Virus treatment of EMT6 tumours altered the tumour infiltrating immune cell proportions where CD8^+^ T cell proportions showed a trending increase over PBS, with VACVΔ4x-SCT^E22^ treatment showing the greatest effect (Figure 6B). There was a trend toward increased proportions of conventional CD4^+^ T cells, and a trend toward decreased proportions of CD4^+^ FOXP3^+^ regulatory T cells in VACVΔ4x-SCT^E22^ treated tumours (Figure 6B). Minimal changes in the proportions of CD19^+^ B cells or CD49b^+^ non-T/non-B cells were observed (Figure 6B). Limited differences in the number of these cell types was also observed (Supplemental Figure 2). Oncolytic virus treatment of EMT6 tumours led to a trending increase in the number of CD45^+^ immune cells recovered from the tumour with the most pronounced increasing trend seen in CD8^+^ T cell numbers (Figure 6C and D). Within the CD8^+^ T cell compartment, there was a non-significant increase in the frequency of CD49b^+^ CD8^+^ T cells of total CD8^+^ T cells in VACVΔ4x-SCT^E22^ treated mice and a trending decrease in the percent of PD-1^+^ CD8^+^ T cells of total CD8^+^ T cells in all oVACV treated groups compared to PBS control (Supplemental Figure 2B). It is of note that CD49b and PD-1 expression were analyzed in a single experimental cohort. To measure tumour antigen specific CD8^+^ T cell responses, we used H-2K^d^ tetramers loaded with the tumour specific E22_(597-605)_ peptide (Figure 6F). We detected a significant increase in the number of E22_(597-605)_ specific CD8^+^ T cells within the tumours of mice treated with VACVΔ4x-SCT^E22^ compared to other virus treatments and PBS control (Figure 6G). Further, VACVΔ4x-SCT^E22^ treatment appeared to induce a systemic anti-tumour response as the total number of E22_(597-605)_ specific CD8^+^ T cells in the spleen was also significantly increased compared to the PBS treated group (Figure 6H). Compared to PBS control, oVACV treatments resulted in an average increase of 2-logs in the number of virus specific CD8^+^ T cells infiltrating the tumour and a 10-fold increase in the spleen, indicating a strong virus directed immune response after treatment (Figure 6G and H) although this increase was only statistically significant in the spleen. In addition, the expression of PD-1 was analyzed on tumour and virus antigen specific CD8^+^ T cells infiltrating the tumour. E22_(597-605)_ specific CD8^+^ T cells showed a significant increase in PD-1 expression after VACVΔ4x-SCT^E22^ treatment compared to non-MHC-I expressing oVACV (Figure 6I). PD-1 expression was also significantly increased on VACVspecific CD8^+^ T cells compared to PBS, regardless of the virus construct (Figure 6I). Collectively, these data demonstrate the induction of both a tumour antigen specific and VACV specific CD8^+^ T cell response intratumorally and systemically when tumour specific pMHC-I is expressed by our novel engineered oVACV.

**Figure 6.**
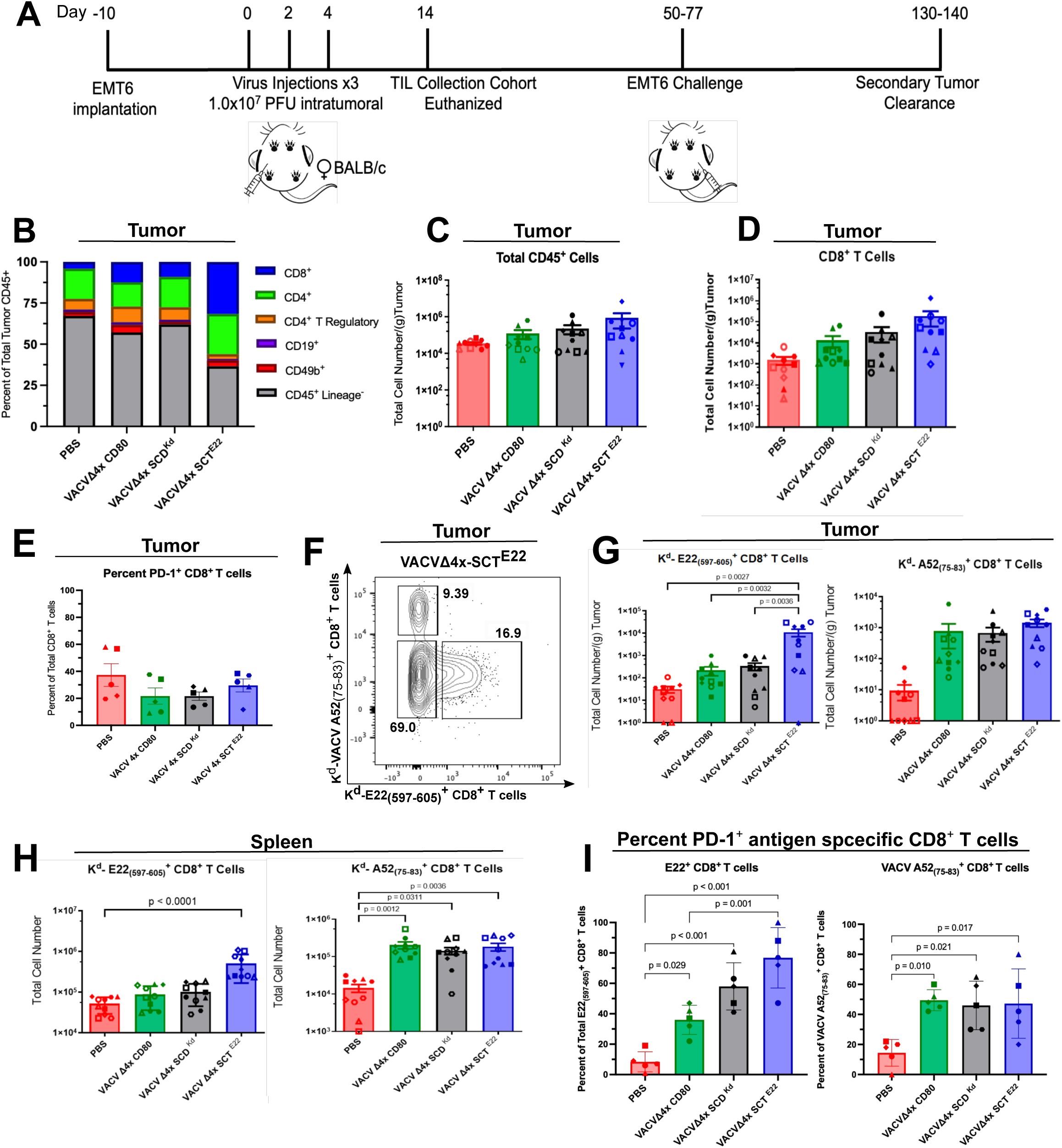
Oncolytic VACV induces altered immune responses and the infiltration of antigen specific CD8^+^ T cells into EMT6 tumours. A, Experimental design to assess survival, tumour growth and tumour infiltrating lymphocyte population changes after treatment with oVACV. Six to eight-week-old female BALB/c mice were injected with tumour cells in the inguinal mammary fat pad at the 4th mammary gland. Once palpable, three intratumoural 1.0×10^7^ PFU doses of oVACVs were administered. A cohort of mice were euthanized 14 days after the first virus dose, where tumour and spleen were collected for immune analysis. B-C, Immune populations and total lymphocytes (CD45^+^) infiltrating the tumour in PBS and oVACV treated mice. D, Total CD8^+^ T cells infiltrating the tumour in PBS and oVACV treated mice. E, Frequency of PD-1^+^ cells of total CD8^+^ T cells infiltrating the tumour in PBS and oVACV treated mice. F, A representative flow cytometry plot for data in G-H is shown. G, E22_(597-605)_ and VACV A52_(75-83)_ antigen specific CD8^+^ T cell numbers infiltrating the tumour from PBS and oVACV treated mice. H, E22_(597-605)_ and VACV A52_(75-83)_ antigen specific CD8^+^ T cell populations in the spleen from PBS and oVACVs treated mice. I, Frequency of PD-1^+^ antigen specific (E22_(597-605)_ and VACV A52_(75-83)_) CD8^+^ T cells infiltrating the tumour in PBS and oVACVs treated mice. Statistical analyses were performed using one-way ANOVA from two independent trials, where p<0.05 is significant and error bars denote mean (SD) or medians and inter-percentile range for data represented on a logarithmic scale.

### Engineered oVACVs enhances EMT6 mammary tumour clearance in a CD8 T cell dependent manner and is improved by anti-PD-L1 checkpoint blockade to induce protective anti-tumour immunity

All PBS treated mice reached endpoint prior to day 50 post first viral treatment (Figure 7A). Virus treatment with VACVΔ4x-CD80 and VACVΔ4x-SCD^Kd^ viruses resulted in some complete responses, 4 out of 17 (23.5%) and 2 out of 17 (11.8%) mice respectively (Figure 7A). However, treatment with VACVΔ4x-SCT^E22^ led to a significant increase in complete responses, where 12 out of 17 (70.6%) mice cleared the tumour (Figure 7A). Since peptide vaccines have also been evaluated as a method to stimulate anti-cancer immune responses,^59^ we wanted to investigate the co-administration of VACVΔ4x-SCD^Kd^ with soluble E22_(597-605)_ peptide as an additional treatment strategy. VACVΔ4x-SCD^Kd^ co-administered with 2 *μ*g E22_(597-605)_ peptide to the tumour did not lead to a difference in the survival of animals treated (1 out of 13) compared to VACVΔ4x-SCD^Kd^ (Figure 7A). To determine if CD8^+^ T cells were necessary for the anti-tumour immune response, anti-CD8α antibodies were administered to deplete CD8^+^ T cells prior to and over the course of oVACV treatment. After CD8^+^ T cell depletion, VACVΔ4x-SCT^E22^ treatment did not lead to tumour clearance in any mice (Figure 7B). We also evaluated whether combining VACVΔ4x-SCT^E22^ and αPD-L1 ICBT could further enhance tumour clearance in the EMT6 model. VACVΔ4x-SCT^E22^ plus αPD-L1 combination therapy improved tumour clearance (8 out of 10 CR) compared to VACVΔ4x-SCT^E22^ plus isotype control treatment (6 out of 10 CR) (Figure 7C), but this difference was not statistically significant. In the EMT6 model, we found that αPD-L1 monotherapy induced some complete responses compared to PBS, but combination treatment with VACVΔ4x-SCT^E22^ and αPD-L1 showed a statistically significant improvement in complete responses. Collectively, these data indicate that oVACV encoding a pMHC-I construct containing a tumour specific antigen significantly improves the efficacy of OV treatment in a model that endogenously expresses MHC-I.

**Figure 7.**
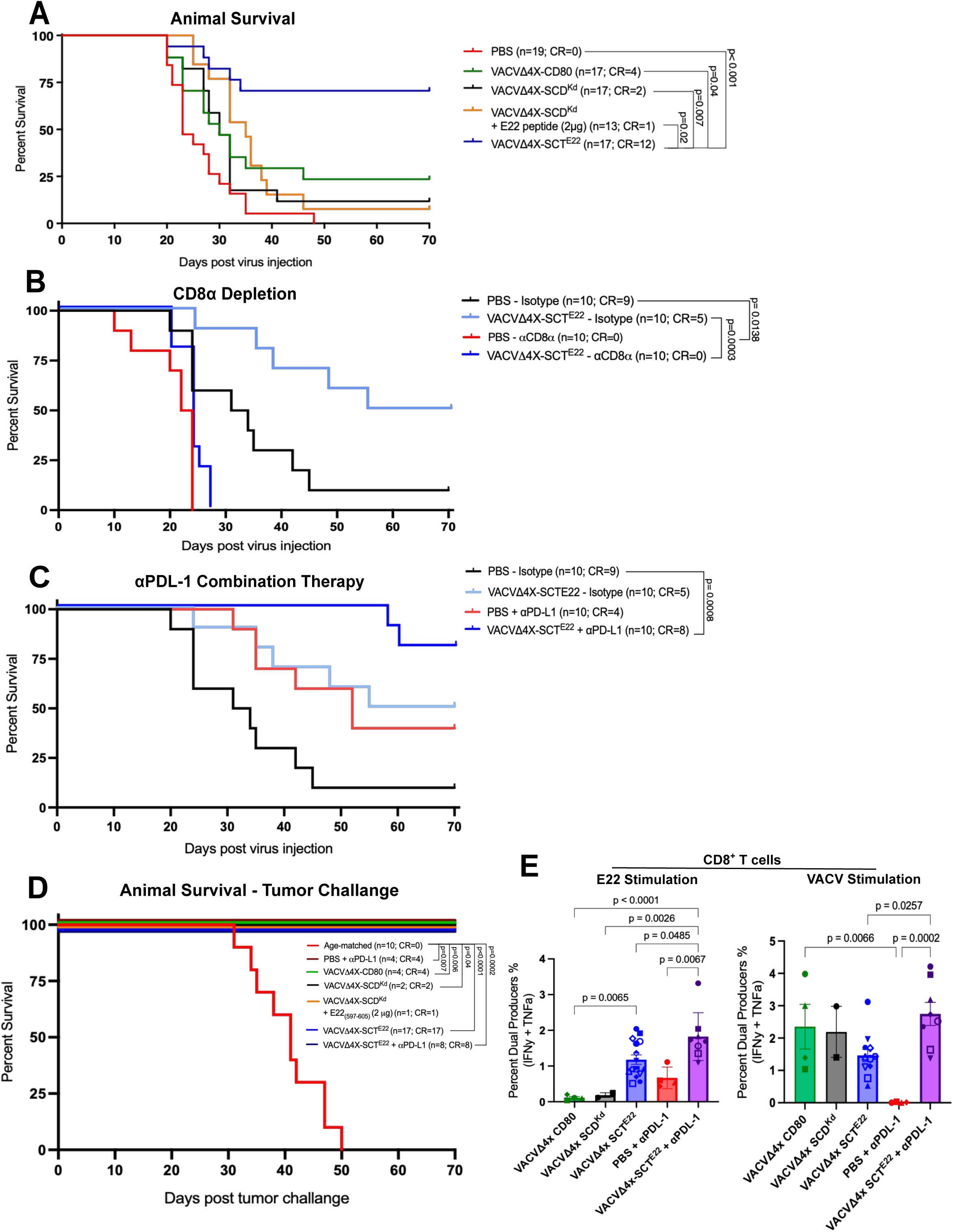
Oncolytic VACVs altered animal survival and therapeutic outcomes in EMT6 tumour bearing mice. A, Kaplan-Meier plot of animal survival after intratumoral treatment with PBS or oVACVs. B, Kaplan-Meier plot of animal survival after intratumoral treatment with PBS or oVACVs during CD8^+^ T cell depletion. C, Kaplan-Meier plot of animal survival after intratumoral PBS or oVACVs treatment and combination αPD-L1 therapy. D, Kaplan-Meier plot of animal survival, after secondary tumour implantation in the opposite inguinal mammary fat pad of primary tumour cleared mice. E, CD8^+^ T cells from mice that cleared primary and secondary tumour produce both IFN*γ* and TNF*⍺* following *ex vivo* peptide stimulation in the presence of Brefeldin A for 4 hours. Statistical analyses were performed using one way ANOVA to compare between groups at day 60 or 70 from two independent experiments, where p<0.05 is significant and error bars denote mean (SD). The represented isotype group data in B-C are the same.

Since treatment of mammary tumours with VACVΔ4x-SCT^E22^ generated robust anti-tumour immune responses in the EMT6 model, we wanted to determine if anti-tumour immunological memory was generated in mice that cleared primary tumour implants following tumour treatment. Rapid tumour rejection was observed in all animals that had cleared primary tumours regardless of the oVACV variant used for treatment (Figure 7D). This indicated a potent anti-tumour immune memory response was induced following primary tumour clearance. In addition to the robust tumour rejection and necessity of CD8^+^ T cells observed in the EMT6 model, we wanted to examine the functionality of the E22_(597-605)_ specific T cells post 2° tumour rejection. Splenocytes from all mice that rejected the 2° tumour were stimulated with either E22_(597-605)_ or a VACV peptide *ex vivo* and the production of IFNγ and TNFα in CD8^+^ T cells was measured. E22_(597-605)_ peptide stimulation resulted in the production of both cytokines and was significantly greater in mice initially treated with VACVΔ4x-SCT^E22^ (Figure 7E). In contrast, animals initially treated with VACVΔ4x-SCD^Kd^ that then cleared 2° tumour injections did not show a robust E22_(597-605)_ specific response (Figure 7E). Enhanced cytokine response to the E22_(597-605)_ peptide was further observed in mice treated with combination VACVΔ4x-SCT^E22^ plus αPD-L1 ICBT. Additionally, mice treated with αPD-L1 monotherapy that cleared both the primary and secondary EMT6 tumour also produced IFNγ and TNFα following E22_(597-605)_ peptide stimulation (Figure 7E), but this was not statistically significant compared to PBS or VACVΔ4x-SCD^Kd^. This supports the ability of ICB therapy to induce or augment tumour specific antigen CD8^+^ T cell responses. Splenic CD8^+^ T cells from all tumour immune mice that were treated with oVACV also produced IFNγ and TNFα in response to VACV peptide regardless of the oVACV variant used for treatment. Overall, these data validate the ability to stimulate tumour antigen specific CD8^+^ T cell responses that are not only capable of inducing tumour cell regression *in vivo* but further results in the generation of robust immunological memory responses that leads to tumour protection upon challenge following primary tumour clearance.

## Discussion

Harnessing the immune system to target and destroy tumour cells underlies some of the most promising cancer therapies. While oncolytic viruses were originally imagined to directly kill cancer cells and eliminate tumours, their efficacy lies in their ability to stimulate an anti-tumour immune response. Modifications to oncolytic viruses to promote anti-tumour immunity have often focused on expanding an antigen agnostic response induced by OV treatment. Here, we showed that a genetically engineered VACV encoding a single chain pMHC-I trimer to focus a CD8^+^ T cell response against a pre-determined tumour antigen and the T cell co-stimulatory molecule CD80 can induce a potent anti-tumour immune response as a curative therapeutic cancer vaccine in pre-clinical models.

Many tumour cells show impairments in MHC-I antigen processing and presentation resulting in substantial reductions in expression of cell surface pMHC-I severely reducing their ability to be recognized by CD8^+^ T cells. We found that infection of B16F10 tumour cells, that are known to have severe defects in MHC-I antigen processing and presentation,^47^ with engineered oVACV expressing an OVA-K^b^ SCT and CD80 resulted in expression of cell surface OVA-K^b^ complexes and CD80 simultaneously. Furthermore, expression of an ‘empty’ MHC-I (SCD) by oVACV in B16F10 followed by pulsing with OVAp resulted in enhanced presentation of pMHC-I complexes. In both instances, activation of OVA-K^b^ specific OT-I T cells *in vitro* was observed (Figure 2E). OVAp-specific CD8^+^ T cells could additionally be expanded *in vivo* when B16F10 tumour-bearing mice were treated intratumourally with VACV expressing the SCT^OVA^ construct (Figure 2F). Therefore, provision of a defined pMHC-I molecule by the virus allows for unimpeded presentation of the peptide of interest, circumventing antigen presentation and processing defects within the tumour cells.

In addition to tumour cell intrinsic impairments in MHC-I antigen presentation reducing the ability of antigen specific CD8^+^ T cells to be activated, OV-mediated induction of an effective anti-tumour CD8^+^ T cell response faces other obstacles. One such obstacle is directing the CD8^+^ T cell response towards tumour antigens given the abundance of viral epitopes that may diminish the amplification of tumour-selective CTLs during infection. While OVs kill tumour cells directly and cause the release of antigens, directing the CD8^+^ T cell response towards a defined tumour antigen is desirable.^60^ For oncolytic VACVs, a variety of strategies have been employed to induce a CD8^+^ T cell response to a given antigen. For example, selected tumour antigens have been integrated into the viral genome for expression by the virus or have been embedded within the envelope of enveloped viruses.^26,61^ However, CD8^+^ T cell responses against many VACV antigens are robust and may dominate the response to any given tumour antigen. While this may not be an issue for highly immunodominant model tumour antigens like OVAp,^62,63^ endogenous tumour associated or tumour specific antigens may not elicit a potent CD8^+^ T cell response. Tethering of the peptide to MHC-I ensures presentation of the TAA and this led to induction of a strong, antigen-specific CD8^+^ T cell response to tumour associated or tumour specific antigens in models that displayed defects in MHC-I presentation or not. Given the ability of APCs to be ‘cross-dressed’ with pre-formed pMHC-I complexes,^64^ our findings support the expression of a such a pre-formed complex by oVACV facilitating antigen presentation and generation of an antigen specific CD8^+^ T cell response.

While the generation of an antigen specific CD8^+^ T cell response to a model foreign antigen, OVA, *in vivo* was demonstrated, a CD8^+^ T cell response to a tumour associated or tumour specific antigen is desirable for tumour clearance and potential to generate anti-tumour immune memory. Intratumoural administration of oVACV expressing a self-antigen from the TAA Trp2 via the SCT^Trp2^ complex to established B16F10 tumours resulted in the expansion of Trp2_(180-188)_ specific CD8^+^ T cells in the tumour and the spleen (Figure 4A and C) and eliminated the tumour in 46% of treated mice. Trp2_(180-188)_ specific CD8^+^ T cells from the tumour expressed the cytotoxic protein granzyme B and were positive for the degranulation marker CD107a and Trp2_(180-188)_ specific CD8^+^ T cells from mice that survived primary B16F10 tumours and rejected secondary B16F10 tumours produced TNFα and IFNγ upon *in vitro* re-stimulation. These findings indicated that self-tolerance to Trp2_(180-188)_ was broken and functional Trp2_(180-188)_ specific CD8^+^ T cells were expanded. Other vaccine approaches including replicating LCMV expressing the Trp2 peptide^65^ and “TriVax” containing peptide, anti-CD40 and CpG^66^ were similarly able to break tolerance and reduce B16F10 tumour growth, however neither treatment appeared to result in tumour clearance. This suggests that engineered oVAC expressing a pMHC-I SCT provided more therapeutic benefit.

Treatment of established EMT6 mammary tumours with oVACV expressing an SCT presenting an EMT6 tumour specific antigen, E22_(597-605)_, resulted in a significant expansion of E22_(597-605)_ specific CD8^+^ T cells in the tumour and spleen, and led to enhanced tumour clearance. As the peptide target chosen was essentially foreign to the host, it is perhaps not surprising that a robust anti-tumour CD8^+^ T cell response was elicited. The effects observed in the EMT6 model also appear to be reliant on the tethering of the peptide to the MHC-I complex, as co-administration of the SCD^Kd^-expressing oVACV and peptide did not yield similar results as the SCT^E22^ treated animals. A stronger effect of peptide administration may have been observed if the peptide was embedded within the virus itself, however, this was not tested. Selection of the peptide may also be important in treatment efficacy. However, a recent clinical study of personalized mRNA-based neoantigen vaccines containing up to twenty MHC-I and MHC-II restricted tumour neoantigens, only generated CD8^+^ or CD4^+^ T cells against their identified tumour neoantigens in half of treated patients, and a further 50% of these responses were restricted to a single neoantigen.^67^

We observed differences in treatment efficacy in the two cancer models. One potential reason for this difference could be due to ability to endogenously express pMHC-I complexes on the surface of the tumour cell. While B16F10 tumours show weak pMHC-I expression, EMT6 tumour cells do not harbour dramatic defects in MHC-I expression. This difference in endogenous pMHC-I expression could allow CD8^+^ T cells specific for EMT6 tumour antigens to exhibit sustained recognition and elimination of tumours while B16F10 tumour recognition may be limited to the period of virus infection.^68^ In addition to the intrinsic differences in MHC-I expression of B16F10 and EMT6 tumour cells, the ‘types’ of peptides presented by the oVACV-delivered pMHC-I complexes is different between the models. Tumour associated antigens are often defined as self-antigens and show evidence of self-tolerance. This creates a barrier to generating a robust CD8^+^ T cell response against these antigens. In contrast, tumour specific antigens, or neo-self-antigens typically display no such intrinsic barrier to generate a CD8^+^ T cell response. The capacity to induce robust anti-tumour responses against a TAA antigen such as Trp2, while possible, are subject to tolerogenic immune pruning unlike neoantigens (E22_(597-605)_) that are not necessarily restricted by immunoregulatory tolerance. While these dramatic differences in the ability to induce CD8^+^ T cell responses exist, it is encouraging that even against longer odds, strong anti-tumour immunity in the B16F10 model was observed.

A variety of immune cell types can be activated and involved in anti-tumour immunity. In the B16F10 model, it was noted that treatment efficacy of VACVΔ4x-CD80 which does not express an MHC-I was not statistically different than treatment with VACVΔ4x-SCT^Trp2^. While CD8^+^ T cells receive the greatest attention, the weak expression of pMHC-I on B16F10 tumour cells may result in NK cell-mediated killing.^69,70^ Early infiltration of the tumour by NK cells has been shown to be required for the recruitment of CD8^+^ T cells. Further, the dynamics of MHC-I expression on B16F10 cells following oVACV administration is unclear and may contribute to the variegated response. A strong IFNγ response has been previously associated with tumour rejection in murine and human studies^71–73^ and has also been shown to induce MHC-I expression on B16F10 *in vitro*.^74^ Since MHC-I expression differentially regulates CD8^+^ T cell and NK cell responses, measuring MHC-I expression on oVACV treated B16F10 *in vivo* should be considered as a future directive for examination. However, the presence of CD8^+^ T cells was required for efficacy of VACVΔ4x-CD80 in B16F10 tumours as CD8^+^ T cell depletion at the time of VACV treatment eliminated any benefit including complete tumour elimination and reductions in tumour growth (Figure 5F). Similarly, CD8^+^ T cell depletion prior to treatment of EMT6 tumours completely abolished any protection provided by oVACV treatment further reinforcing the critical role played by CD8^+^ T cells in oVACV therapy (Figure 7F).

Given the requirement for CD8^+^ T cells in promoting tumour clearance in both the B16F10 and EMT6 models, it is perhaps not surprising that combination therapy with αPD-L1 checkpoint blockade enhanced complete responses in both models. As we showed, αPD-L1 treatment had no benefit as a monotherapy in the B16F10 model, but significantly enhanced responses with both VACVΔ4x-CD80 and VACVΔ4x-SCT^Trp2^ treatment. Immune checkpoint blockade monotherapy showed some efficacy in eliminating EMT6 tumours, though combination therapy with VACVΔ4x-SCT^E22^ showed enhanced responses over either monotherapy. VACVΔ4x-SCT^E22^ plus αPD-L1 combination treatment additionally enhanced the production of cytokines in CD8^+^ T cells isolated from the spleens of animals that cleared 1° and 2° tumours when stimulated with soluble E22_(597-605)_ peptide. Interestingly, combination therapy did not similarly enhance the response to the Trp2_(180-188)_ peptide in the B16F10 model suggesting that ICBT may augment responses to other CD8^+^ T cell epitopes through epitope spreading.^75^ Together, this demonstrates the ability to increase the potency of the immune response that is induced when combination ICB therapy is used with oVACV.

One important outcome of inducing a robust anti-tumour immune response is the ability to establish anti-tumour immune memory. Should immune memory be generated following treatment and elimination of tumours, growth of future tumours can be prevented. Of the mice that cleared a B16F10 tumour following oncolytic VACVΔ4x-CD80 treatment, only 50% of those cured mice rejected a secondary B16F10 tumour implanted at least 30 days later. Further, this limited memory response was not enhanced when VACVΔ4x-CD80 treatment was used in combination with αPD-L1. In contrast, most of animals rejected 2° tumours when they were initially treated with VACVΔ4x-SCT^Trp2^ with or without ICBT, indicating the importance of expressing a TAA during primary treatment to generate a potent memory immune response. This aligns with the presence of Trp2_(180-188)_ specific CD8^+^ T cells that respond to Trp2_(180-188)_ peptide in mice that cleared primary B16F10 tumour and survived tumour rechallenge. On the other hand, mice cured of primary EMT6 tumours, regardless of the therapy, universally rejected a secondary EMT6 tumour. Furthermore, we detected a robust, peptide specific CD8^+^ T cell response from the mice that cleared a primary and secondary EMT6 tumour, indicating a strong CD8^+^ T cell memory population was present.

Collectively, this work demonstrated the utility of a VACV based oncolytic virus expressing a defined tumour specific peptide-MHC-I complex to treat mice with established tumours.

However, there are a number of limitations to this approach for clinical translation. First, identifying appropriate tumour associated or tumour specific antigens in combination with HLA-typing is needed. Relatively straightforward protocols have been developed in the transplantation area and can be employed in this regard. Genomic sequencing of tumours can be used to identify potential antigens and algorithms to predict antigen presentation are being advanced.^76,77^ Finally, while not a focus of this study, determining an optimum route of administration is important for clinical translation. Currently, intratumoural administration of oncolytic viruses is utilized clinically, but the use of other routes may be more desirable provided clinical efficacy is maintained. Moving forward, this approach provides a compelling therapeutic opportunity that could yield either an off-the-shelf or personalized treatment option.

## Materials and Methods

### Cell lines and Culture Conditions

B16F10 mouse melanoma cells (ATCC CRL-6475) were cultured in DMEM containing 2 mM L-glutamine, 100 U/mL antimycotic antibiotic, 1X nonessential amino acids, 1 mM sodium pyruvate, 1 µM β-mercaptoethanol, supplemented with 10% heat inactivated Fetal Bovine Serum (HI-FBS). The EMT6 murine mammary carcinoma cell (provided by Dr. Mary Hitt University of Alberta, ATCC CRL-2755) were cultured in RPMI-1640 containing 100 U/mL penicillin, 0.1 mg/mL streptomycin supplemented with 10% HI-FBS (Gibco). BSC-40 cells (ATCC RRID: CVCV_3657), received in 2005 from ATCC were cultured in MEM supplemented with 5% FetalGro bovine growth serum (RMBIO). Cells were passaged for fewer than 16 weeks and tested for mycoplasma contamination using a Mycoplasma PCR Detection Kit (Sigma Milipore).

### Viruses

All viruses were constructed from a clonal isolate of Vaccinia Virus (VACV) strain Western Reserve using traditional homologous recombination techniques (37). The parental VACVΔ3x (Δ*J2R*, Δ*B8R*, Δ*B18R*) was previously described (38). The additional removal of the *M2L* locus (39) resulted in the generation of the VACVΔ4x (Δ*J2R*, Δ*M2L*, Δ*B8R*, Δ*B18R*) viral backbone used to develop all variants discussed herein. To generate the remaining viruses, we used the pUC57 plasmid encoding one of three genes; the SCT construct; a peptide covalently joined to the β2microglobulin via a flexible linker further covalently joined to the major histocompatibility complex (MHC) heavy chain by a flexible linker as a single chain protein, the SCD construct; β2microglobulin covalently joined to the major histocompatibility complex (MHC) heavy chain by a flexible linker, or the CD80 protein. Each gene was flanked by regions of homology to the target locus allowing for site specific recombination. BSC-40 cells were infected with VACVΔ4x at a MOI of 0.1 and transfected two hours post infection with 2.5 μg of linearized plasmid DNA using Lipofectamine 200 (Invitrogen). Progeny viruses were collected 24 hours post transfection and purified using two rounds of drug selection followed by up to four rounds of plaque picking under agar. PCR was used to verify incorporation of the virus modifications (Table 1). Virus stocks were produced by infection of BSC-40 cells for 72 hours, and the harvested cells were lysed using Dounce homogenization as previously described (38). Virus preparations were purified on a 36% sucrose cushion and virus titres determined by plaque assay on BSC-40 cells. Full virus genome sequencing was performed by Plasmidsaurus using Oxford Nanopore Technology or Illumina technologies to confirm virus fidelity.

### Virus Characterization

Multistep growth curves and cytotoxicity assays were performed as previously described (38). Briefly, BSC-40 cells at 80% confluency were infected with oVACV variants at a MOI of 0.03 for 1 hour, after which fresh media was added on top and the plate was returned to 37°C. At multiple time points post-infection (0, 3, 6, 12, 24, 48, 72 hours) cell monolayers and supernatant were collected by cell scraping and freeze-thawed three times at -80°C to lyse cells. Cell lysates were diluted, plated in triplicate on BSC-40 cells, and incubated for 48 hours at 37°C in media containing 1% carboxy-methylcellulose (Sigma). Cells were subsequently fixed and stained with crystal violet for plaque counting to determine viral titres.

Cytotoxicity assays were performed in 96-well plates by seeding two thousand BSC-40 cells/well and infecting with the various virus constructs 24 hours post seeding. Seventy-two hours post infection the cell culture media was replaced with fresh media containing 44 μmol/L resazurin (ThermoFisher) and plates were incubated for 3 hours at 37°C. Fluorescence was measured using a microplate reader (Promega) at 560 ηm excitation and 590 ηm emission.

### *In vitro* T cell Activation Assays

B16F10 tumour cells were infected with oVACV variants at a MOI of 3 for 7 hours at 37°C. Splenocytes from a TCR transgenic OT-I mouse were harvested the day of viral infection and labelled with cell proliferation dye. Briefly, 1 μL of 5 μM carboxyfluoroscein succinimidyl ester (CFSE) was added to 1 mL of 1.0×10^7^ cells/mL in PBS and incubated for 10 min at 37°C with intermittent mixing by inversion. A 5X volume of serum containing media was added, and the cell suspension incubated for 5 min at room temperature to quench the CFSE. CFSE-labelled cells were washed once and resuspended in sterile RP10 media (RPMI 1640 with 2.05 mM L-glutamine [HyClone], 10% HI-FBS, 5 mM HEPES, 50 μM 2-ME, 50 mg/mL penicillin/streptomycin). Seven hours after infection, the infected tumour cells were harvested using cell scraping and a single cell suspension was mixed with CFSE-labelled OT-I T cells at a 1:1 ratio in a 48 well plate and incubated at 37°C. The co-cultures were collected at the indicated time points, transferred to Eppendorf tubes and washed with PBS prior to processing for flow cytometry as described below.

### *In vivo* animal models

Animal studies were carried out at the University of Alberta (Edmonton, Alberta, Canada) and completed in accordance with the Canadian Council on Animal Care Guidelines and Policies with approval from the Animal Care and Use Committee: Health Sciences for the University of Alberta.

Male C57BL/6 and female BALB/c mice between 8-12 weeks old were purchased from Charles River. Animals were housed in high-efficiency particulate air (HEPA)-filtered ventilated cages in a biosafety level 2 containment suite at the University of Alberta North Campus Animal Services facility in groups between three and five mice and given at least 1 week to acclimate to the housing facility. Environmental conditions included a temperature of 21°C ± 2°C, humidity of 55% ± 10%, lighting of 350 lux, and a 12:12 light:dark cycle with lights on at 07:00 and off at 19:00. Animals were housed in 595 x 380 x 200 mm cages (Ehret) and given access to mouse maintenance food (LabDiets) and water freely. Environmental enrichments including bedding, one red tinted mouse tunnel (Bio-Serv) one 50 mm x 50 mm Nestlet (Bed’r’Nest) and one 4-8 g portion-controlled nesting material (Ancare) were provided.

For orthotopic tumours, tumour cells were harvested by trypsinization, washed twice and resuspended in PBS at the desired concentration. Prior to injection, 25 μL of Matrigel (Corning) was combined with 25 μL of cells in PBS containing 1.0×10^5^ B16F10 or EMT6 cells per mouse.

The Matrigel-cell mixture was injected subcutaneously in the flank of male mice for the B16F10 model or into the inguinal mammary fat pad of female mice below the fourth nipple for the EMT6 model. Tumours were palpated to monitor growth and were measurable using calipers approximately 7-8 days after implantation for the B16F10 tumour model and 9-10 days in the EMT6 model allowing tumours to reach an average volume between 14-30 mm^3^ and 15-70 mm^3^ respectively at the time of the first virus injection.

Individual mice were randomized into treatment groups. Viruses were prepared for animal use by sonicating and diluting virus stocks to 2.0×10^8^ plaque forming units (PFU)/mL in filtered PBS.

For virus injections, mice were anesthetized with isoflurane and injected intratumourally with 50 μL of virus yielding a dose of 1.0×10^7^ PFU per injection. Two additional doses were administered at 48-hour intervals. Tumour growth was measured using calipers twice weekly and tumour volumes calculated with a modified ellipsoid equation: = (1/24) x L x (W+H)^2^. Mice were euthanized by CO_2_ inhalation at a tumour burden endpoint of 1,500mm^3^ or at a clinical endpoint defined by a combination of scored indicators (ulcerated tumour, hunched posture, ruffled fur, unresponsiveness to touch or weight loss exceeding 15% of body weight). Mice that cleared the tumour and remained tumour free for 30 days were challenged with B16F10 or EMT6 cells parallel to the primary tumour on the opposite flank or inguinal mammary fat pad, respectively. Secondary implanted tumours were monitored for growth as described above.

Age matched control mice, naïve to tumour and virus treatment, were implanted with tumour cells to validate observations in the challenged mice. Mice were categorized as having cleared tumour at 30 days post euthanasia of the final age matched control using the endpoint described above.

### Tissue Processing

Spleens were mashed between two 70 µm mesh squares into splenocyte isolation buffer (PBS + 2% HI-FBS + 0.5 mmol/L EDTA) using the rubber end of a 3 mL syringe. The isolated cells were centrifuged at 400xg for 5 min and resuspended in 2 mL of 1X ACK (Ammonium-ChloridePotassium) lysis buffer (distilled H_2_O + 0.15 M ammonium chloride + 10 mM potassium bicarbonate + 0.1 mM disodium EDTA) for 5 min. Cells were returned to isotonicity by the addition of 10 mL isolation buffer, centrifuged at 400xg for 5 min and washed twice with isolation buffer.

Tumours were harvested and collected into 5 mL of RPMI-1640 containing 10% FBS, 0.5 mg/mL collagenase type 1A (Sigma-Aldrich) and 10 µg/mL DNase I (Roche). Tumours were cut into small pieces using scissors before being dissociated using the m_impTumour01.01 protocol on a GentleMACs dissociator (Miltenyi Biotec). The tumour samples were then incubated with shaking for 30 min at 37°C. After dissociation and shaking, the tumour chunks were filtered through a 70 µm cell strainer into PBS and centrifuged at 400xg for 5 min. For B16F10 tumours, infiltrating leukocytes were enriched from the tumour cells on a 40%/80% Percoll gradient (Sigma-Aldrich). The gradient was centrifuged at 300xg for 30 min at room temperature, leukocytes concentrated at the 40%/80% interface were collected, washed with PBS, and resuspended in sterile RP10 media prior to flow cytometry staining.

### Flow Cytometry Staining

2.0×10^6^ splenocytes or cell co-cultures were transferred to individual wells of 96-well round bottom plates. TIL recovered from tumours were resuspended in 200 μL of media and transferred to a 96-well plate for staining. For control PBS treated tumours, single cell suspensions were resuspended in ∼500 μL of media and 200 μL was used for staining. Cells were washed with PBS and stained with blue-fluorescent reactive dye (Invitrogen) for 20 min at room temperature. Surface antibody staining was then completed using the antibodies described in Table 2 at 4°C for 30 min, and fixed and permeabilized (BD Bioscience) for intracellular staining. Vaccinia antibodies were produced in-house as previously described (38) and added at 1:1000 for 1 hour at 4°C, washed twice and stained with Alexa Fluor 647 fluorescently labelled goat anti-mouse secondary antibody at 1:2000 for 30 min at 4°C. Samples were run on the BD LSR Fortessa or Cytek Aurora Flow Cytometer and analyzed with FlowJo Software (TreeStar). A representative example of the cell-gating strategy is found in Supplementary Figure 1.

### Intracellular Cytokine Staining

Spleens were collected from mice that cleared both primary and secondary tumour implantations, processed as above and stimulated with 1.0×10^−5^ mg/mL soluble peptide *in vitro* for 4 hours at 37°C. Peptide specific to the tumour model (Trp2_(180-188)_ in B16F10 or E22_(597-605)_ in EMT6) or VACV peptide pools (A8_(189-96),_ A23_(297-305),_ A47_(138-146),_ A47_(171-180),_ and A52_(75-83)_) were used. Brefeldin A (1000X) was added to stimulated cells at 1 μL/mL to sequester cytokine production within cells for the duration of the stimulation period. Intracellular cytokine staining was performed as described above.

### Statistical Analysis

Data were analyzed using GraphPad Prism 9. If data were determined to be normally distributed by the Shapiro-Wilk normality, parametric one-way ANOVA testing was performed with Tukey’s multiple comparisons. Significance was set at p ≤ 0.05. Virus growth curves were analyzed using a two-way ANOVA and Tukey’s multiple comparisons test to compare virus titre at each time point. Survival data were analyzed using simple survival analysis, Kaplan-Meier and significance by log-rank (Mantel-Cox) with multiple comparisons using the Holm-Šídák method. For *in vivo* studies, the significance threshold was adjusted for multiple comparisons, where the significance threshold was set as p=0.05/K where K is the number of comparisons being made.

## Acknowledgements

Flow cytometry experiments were performed at The University of Alberta Faculty of Medicine & Dentistry Flow Cytometry Facility, which receives financial support from the Faculty of Medicine & Dentistry and Canada Foundation for Innovation (CFI) awards to contributing investigators. Health Sciences Laboratory Animal Services provided animal husbandry and technical assistance.

This work was supported by grants from the Mary Johnston Family Melanoma Research Program and Alberta Cancer Foundation (to T.A.B), the Li Ka Shing Institute of Virology (to T.A.B, R.S.N and D.H.E), the Canadian Institutes of Health Research (to T.A.B) and Striving for Pandemic Preparedness - an Alberta Research Consortium (to T.A.B and R.S.N). Studentship support was received from the Li Ka Shing Institute of Virology, the Faculty of Medicine and Dentistry (to S.K. and J.W), and the Canadian Institutes of Health Research (to J.W.).

T.A.B, R.S.N and D.H.E conducted conceptualization, funding acquisition and project administration. S.K., J.W. and N.F. designed the methodology, conducted the investigation, data curation and formal analysis for the experiments. A.C. contributed experiment execution and data curation. S.K. wrote the paper with significant contributes by T.A.B, R.S.N and S.W.

