## Supplemental Tables and Figures for "Oncolytic vaccinia virus expression of a defined peptide-MHCI complex as a precision cancer immunotherapy platform"

Table S1. PCR primers for assessing viral purity.

| Target | Primer Set | Wildtype Amplicon size | Mutant Amplicon size |
|---|---|---|---|
| J2R | 5' TATTCAGTTGATAATCGGCCCCATGTTT<br>5' GAGTCGATGTAACACTTTCTACACACCG | 516 bp | 844 bp |
| M2L | 5' TCCCGACATTAAATTGGCTATAG<br>5'CCGTCTCATTTGGAGAGTATC | 1328 bp | 2342 bp |
| B8R | 5' ATCACTTCAGTGACAGTAGTC<br>5' AGGACTATAATCAGGGACCTC | 966 bp | 1067 bp |
| B18R | 5' CCACCTACCAAAGTATAGTTG<br>5' CGGTGAGATACAAATACCTAG | 1250 bp | 269 bp |

Table S2. Flow cytometry antibodies used in the current study.

| Antibody | Clone | Company |
|---|---|---|
| CD4 | RM4-4 | Biolegend |
| CD8 | 53-6.7 | BD Biosciences |
| CD45.2 | 104 | Invitrogen |
| CD19 | 1D3/CD19 | Biolegend |
| CD44 | IM7 | Invitrogen |
| CD25 | PC61 | Biolegend |
| CD69 | H1.2F3 | Biolegend |
| PD-1 | 29F.1A12 | Biolegend |
| CD49b | DX5 | Biolegend |
| NK1.1 | PK136 | Biolegend |
| FoxP3 | FJK-16s | Invitrogen |
| CD107a | 1D4B | Biolegend |
| Granzyme B | GB12 | Invitrogen |
| IFN$\gamma$ | XMG1.2 | Invitrogen |
| TNF$\alpha$ | MP6-XT22 | Invitrogen |
| H-2K$^b$ bound to OVA$_{(257-264)}$ | 25-D1.16 | Biolegend |
| H-2K$^b$ OVA$_{(257-264)}$ SIINFEKL – PE Tetramer | Not applicable | NIH Tetramer Facility |
| H-2K$^b$ Trp2$_{(180-188)}$ SVYDFFVWL – BV421 Tetramer | Not applicable | NIH Tetramer Facility |
| H-2K$^d$ VACV A52$_{(75-83)}$ KYGRLFNEI – BV421 Tetramer | Not applicable | NIH Tetramer Facility |
| H-2K$^d$ E22$_{(75-83)}$ RYAQAFTLL – PE Tetramer | Not applicable | NIH Tetramer Facility |

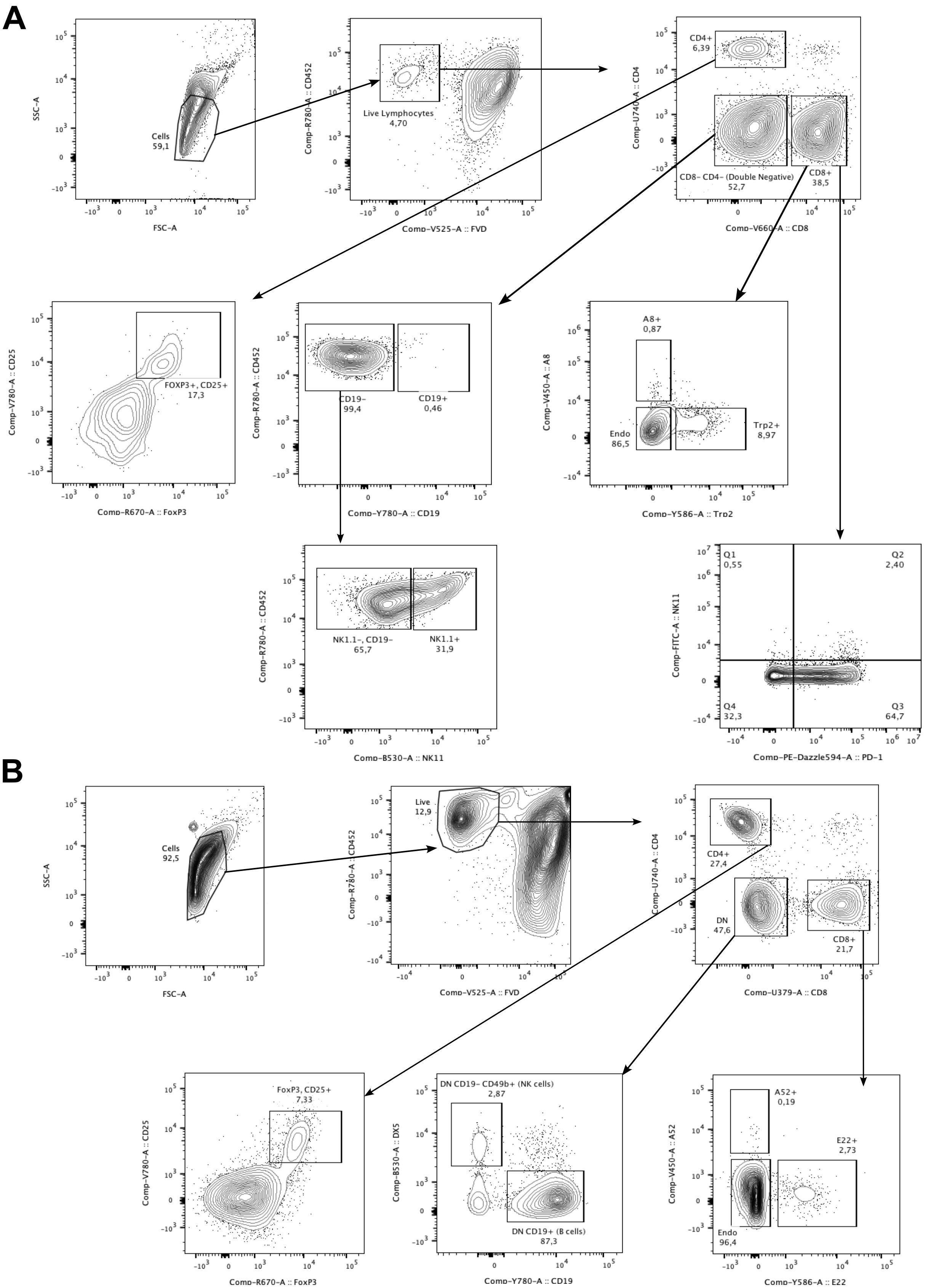


Figure S1. Ṙepresentative gating strategy for identifying immune cell subsets in tumour and spleen samples. A. B16F10 representative spleen gating. Briefly, splencoytes were gated based on FSC-A and SSC-A and CD45.2 expression. Dead cells were removed from analysis based on uptake of viability dye Aqua Zombie. T cells were identified by $CD4^+$ and $CD8^+$ marker expression. T regulatory cells were isolated from the $CD4^+$ subset, further gated on $FOXP3^+$ $CD25^+$ populations. Antigen specific T cells were gated on the $CD8^+$ population with tetramer staining. Each subset was the further gated on PD-1, and NK1.1 markers. Other immune populations including B cells were gated from double negative CD4- and CD8- gates, where B cells are CD19+ and NK-like cells are CD19- and $NK1.1^+$ or $CD49b^+$. B. EMT6 representative spleen gating, groups identified using the same path as B16F10.

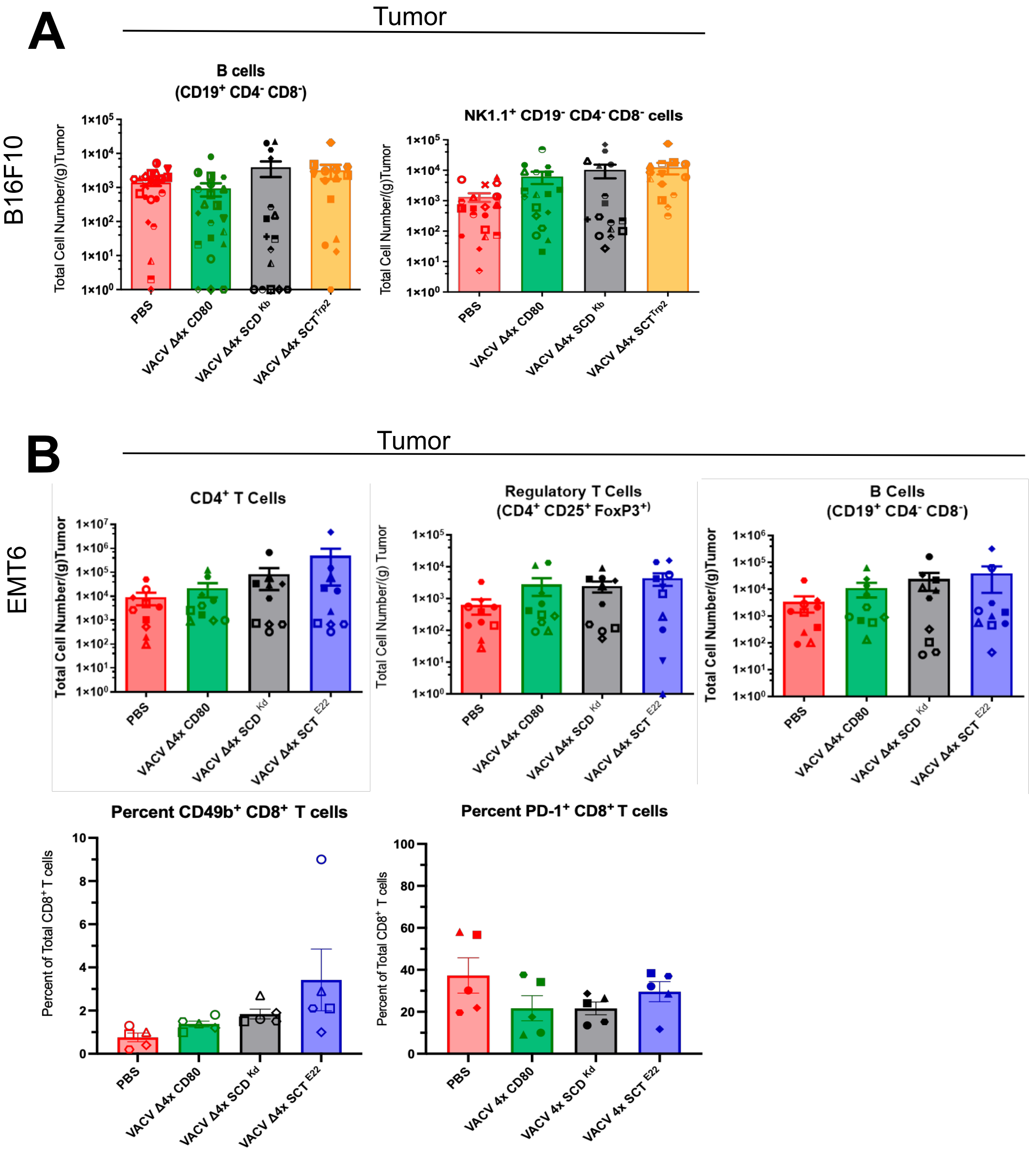


Figure S2. Additional immune cell populations infiltrating the tumor.

A. Additional immune cell subset infiltrates isolated from B16F10 tumours treated with PBS or oVACVs. Minimal differences are observed with B cell and natural killer-like cell numbers. One way ANOVA was used to evaluate differences in total cell number/(g) tumour between groups at day 14 from two independent experiments. B, Additional immune cell subset infiltrates from EMT6 tumours treated with PBS or oVACVs. $CD4^+$, $CD8^+$ and regulatory T cell numbers increased with oVACV treatment, where minimal differences are observed with B cell numbers. Natural killer-like cells trend toward an increase only following VACVΔ4x-$SCT^{E22}$, this marker was analyzed in one experiment, n=5. One way ANOVA was used to evaluate differences in total cell number/(g) tumour between groups at day 14 from two independent experiments, where $p<0.05$ is significant.

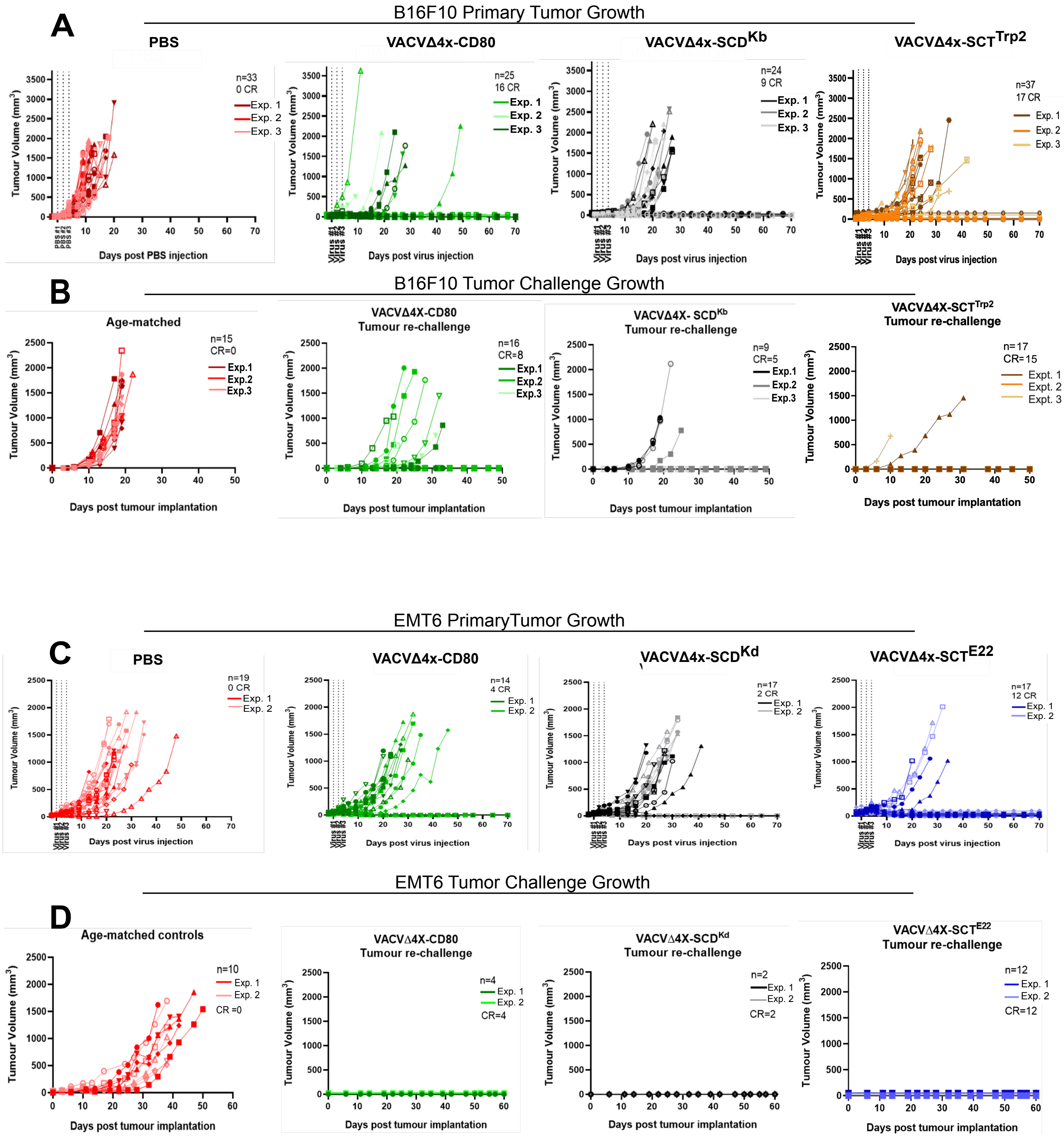


Figure S3. Tumor growth after primary implantation, treatment, and challenge.
A, Tumour growth and regression after primary tumor implantation and treatment with PBS or oVACVs in the B16F10 model. CR, complete response with no tumour for 30 days post initial clearance. B, Tumour growth and regression after secondary tumour implantation in the opposite flank of primary tumour cleared mice in the B16F10 model. CR, complete response achieved as mice reject implants and remain tumour free for 30 days. C, Tumour growth and regression after primary tumor implantation and treatment with PBS or oVACVs, in the EMT6 model. CR, complete response with no tumour for 30 days post initial clearance. D, Tumour rejection after secondary implantation in the opposite mammary fat pad of primary tumor cleared mice in the EMT6 model. CR, complete response following tumour rejection and remained tumour free for 30 days.
